# The Human Cell Line Phosphoproteome Atlas: A Deep Empirical Resource Revealing Kinase Activity Landscapes

**DOI:** 10.64898/2025.12.05.692571

**Authors:** Claire Koenig, Hayoung Cho, Kristina B. Emdal, Ilaria Piga, Pierre Sabatier, Samuel Lozano-Juárez, Ana Martinez-Val, Jesper V. Olsen

## Abstract

Protein phosphorylation orchestrates cellular signaling and controls diverse biological processes, with its dysregulation driving diseases, notably cancer. Comprehensive, high-throughput phosphoproteomics remains limited by detection sensitivity, data completeness, and computational bottlenecks, especially in low-input settings. We present the deepest empirical human phosphoproteome resource to date, regrouping over 200,000 class I phosphosites across 33 diverse human cell lines, and demonstrate that this spectral library dramatically improves single-shot phosphoproteomics with 30-fold faster data processing compared to library-free approaches and enhanced confidence in phosphosite localization even from minimal sample input. Integrating proteome and phosphoproteome data, we developed a combined kinase activity score, revealing cell line- and cancer-specific signaling vulnerabilities, many correlating with drug sensitivity. This resource accelerates deep, reproducible phosphoproteomics, enabling systematic functional mapping of cellular signaling networks, and empowers precision oncology by highlighting actionable kinase targets in diverse cell states.

## Introduction

Cellular signaling is dynamically regulated by thousands of coordinated site-specific protein phosphorylation events that determine how cells grow, respond to stimuli, or promote disease. Protein phosphorylation is catalyzed by protein kinases and erased by protein phosphatases, and the cellular interplay between these enzymes are usually tightly controlled, making protein phosphorylation one of the most important post-translational modifications (PTMs). Phosphorylation-dependent mechanisms control diverse biological processes, and each cell type, with its unique function and regulatory circuitry, is expected to exhibit a distinct phosphoproteome. Beyond enhancing our understanding of cellular biology, global phosphoproteomics has demonstrated efficiency in understanding cancer biology, where dysregulated kinase signaling rewires essential communication networks and promotes aberrant cellular progression, providing actionable insights into therapeutic vulnerabilities ^1–3^. Mass-spectrometry (MS)-based phosphoproteomics has become the primary method for simultaneously identifying and quantifying protein phosphorylation events, with up to ∼119,000 phosphosites detected in large-scale studies ^4,5^, providing invaluable insights into the regulatory layers governing cellular behavior ^6,7^. However, extracting meaningful biological information requires datasets that are as complete and comprehensive as possible. While large-scale initiatives aimed at making large genomics, transcriptomics and proteomics datasets available across tissues or cancers, such as the Human Protein Atlas ^8^, the Dependency Map ^9,10^ or CPTAC ^11^, phosphoproteomics profiling is often absent, or limited to cancer material.

Despite rapid and substantial developments in the field of phosphoproteomics, accessing the full depth of the phosphoproteome remains challenging. Many phosphorylation events are highly dynamic and occur at very low stoichiometry, spanning across a wide abundance range, making their detection by liquid chromatography tandem mass spectrometry (LC-MS/MS) challenging. Experimental MS-based phosphoproteomic workflows depend on enrichment strategies that may bias the detectable phosphosite populations, and MS acquisition can only sample peptides within specific mass over charge and intensity windows, inherently limiting phosphoproteome coverage. Historically, quantifying 30,000 phosphosites required large input amounts and extensive offline fractionation ^12^. Today, technical advances combining fast and sensitive LC-MS/MS have enabled comparable coverage in single-shot analyses with dramatically less material, including reports of ∼ 30,000 phosphosites measured in 30-minute gradients ^13,14^. Moreover, downscaled workflows have demonstrated remarkable sensitivity using minimal input amounts as low as tens of nanograms ^15–17^ , bringing phosphoproteomics closer to feasibility for clinical applications. Unlike conventional bottom-up proteomics in which most proteins are robustly identified and quantified by many peptide precursors, phosphoproteomics mainly relies on single precursor-level quantification and confident site localization, which greatly increases susceptibility to missing values, particularly in low-input settings.

While data-dependent acquisition (DDA) was historically the method of choice for phosphoproteomics ^18,19^, the advent of spectral-library-free data-independent acquisition (DIA) approaches have simplified experimental design and substantially increased coverage ^20^. Yet, the rising scale of modern studies with automated sample processing workflows, combined with the high acquisition speed of modern MS instruments, creates a new bottleneck: searching and processing the resulting massive LC-MS/MS datasets, which demands significant computational resources and long search times.

Here, we leveraged state-of-the-art narrow-window DIA (nDIA) analysis ^21^ to investigate the extent and regulation of the MS-detectable human phosphoproteome across a panel of 33 human cell lines representing the major organs, tissues and cancer cells in the body. Where previous efforts reported up to 119,000 phosphosites ^4,5^, our goal was to increase the depth further and build a comprehensive empirical spectral library of MS-detectable human phosphopeptides. In parallel, we evaluated whether the large empirically-generated phosphopeptide library can reduce the computational load and accelerate phosphoproteomics data processing without compromising analytical depth. Finally, we take advantage of the depth of data generated to uncover distinct signaling mechanisms across the panel of human cell lines. Based on this, we developed a kinase activity score that revealed cell-type specific kinase vulnerabilities that explained cell-line specific kinase inhibitor drug sensitivity.

### I. Analysis of 33 human cell lines results in exceptionally high coverage of the human proteome and phosphoproteome

#### Building of the human cell lines panel and generation of the proteomics and phosphoproteomics profiles

To create a uniform and comprehensive experimental human phosphopeptide library representing phospho-signaling in living cells, we assembled a panel of 33 different human cell lines, which were selected to capture lineage diversity and represent the major cell types and signaling pathways in the human body (**Figure 1A**). The panel included both cancerous and non-cancerous phenotypes, comprising immortalized cancer cell lines as well as primary cell lines. To ensure broad representation, we included suspension and adherent cell lines, including epithelial, mesenchymal, endothelial, and fibroblast morphologies, while early developmental signaling was covered by incorporating embryonal cells and induced pluripotent stem cells. The cell lines were profiled in both unstimulated serum-starved and serum-stimulated conditions, the latter inducing a broad range of regulatory phosphorylation events. Additionally, increased phosphotyrosine coverage was achieved by broad-spectrum inhibition of phosphotyrosine protein phosphatases via pervanadate treatment ^22,23^ of the colorectal cancer cell line DLD-1. Four biological replicates of both unstimulated and serum-stimulated conditions, were collected for each cell line to identify statistically significantly serum-regulated phosphosites..

**Figure 1:**
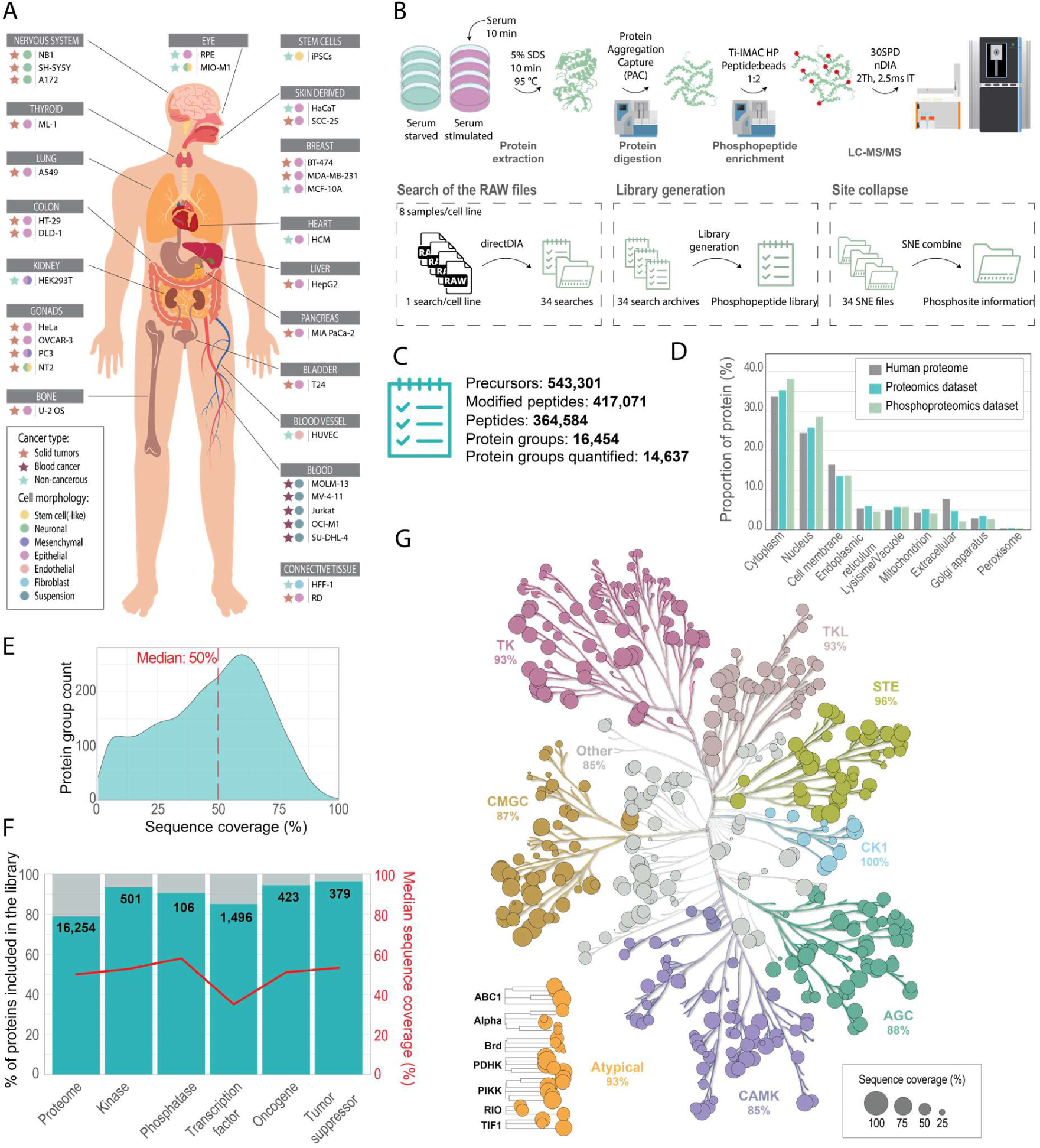
Overview of the cell line panel, experimental workflow, and proteome coverage. A-Overview of the 33 human cell lines used to build the panel, organized according to their tissue of origin. B- Overview of the experimental workflow applied for the generation of the data set. Cells were cultured, and half of the replicates were serum-stimulated before lysis, followed by protein digestion, phosphopeptide enrichment, and LC-MS/MS analysis. The generated raw files were searched per cell line and combined for the generation of (phospho)peptide libraries. C- Overview of the depth of the resulting peptide spectral library. D- Distribution of the cellular localization of the proteins using predicted localization with DeepLoc ^28^ in the human proteome, and the generated proteomics and phosphoproteomics datasets. E- Protein sequence coverage distribution over the 16,454 protein groups. F- Barplot representing the percentage of proteins included in the peptide library over various protein families. The red line represents their respective median protein sequence coverage. G- Kinase map generated using KinMap ^25^, where each dot on the kinase tree represents a kinase in the peptide library and the size of the dot represents the protein sequence coverage. The percentages on the plot display the fraction of kinases detected in the library for a given kinase family.

All cell lines were harvested in boiling SDS-buffer to instantaneously quench all enzymatic activity and thereby preserve the cellular phosphorylation state at the time of lysis, during sample preparation and MS analysis. Whole cell lysates were processed by magnetic bead-based protein aggregation capture (PAC) and on-bead digested with LysC and trypsin. The resulting tryptic peptide mixtures were enriched for phosphopeptides via magnetic bead-based zirconium ion immobilized metal affinity chromatography (Zr-IMAC) prior to label-free LC-MS/MS analysis. Both unenriched peptide mixtures representing the cell line proteomes and phosphopeptide-enriched samples were analyzed using an Evosep One LC system coupled to an Orbitrap Astral mass spectrometer operated in nDIA mode (**Figure 1B**). Each cell line was analyzed individually using Spectronaut (v19) by searching the eight replicates of serum-stimulated and unstimulated samples together. Peptide and phosphopeptide libraries were generated by combining the corresponding search archives while maintaining false-discovery rate (FDR) control and ensuring homogeneous protein inference. For downstream analysis, the Spectronaut experiment (SNE) files were merged and the associated quantitative tables were exported (**Figure 1B**).

#### Deep proteome library covers up to 80% of the canonical human proteome

Deep human cell line proteome coverage was achieved by using a 30 samples-per-day (SPD) 44-minute LC gradient with the fastest scanning 200 Hz nDIA MS acquisition method. This resulted in an average quantification of 9,790 protein groups per file, with a maximum of 10,616 and a minimum of 8,966 across cell lines (**Suppl. Fig. 1A**). Collectively, this resulted in the generation of a spectral library comprising 543,301 peptide precursors, corresponding to 364,584 unique peptide sequences and 16,454 protein groups, of which 14,637 remained after filtering for quantified protein groups in at least three replicates in at least one cell line (**Figure 1C, Suppl. Fig. 2**). The quantified proteome was predominantly localized to the cytoplasm and the nucleus, as assessed by DeepLoc predictions ^24^ (**Figure 1D**), with a distribution across cell compartments following the expected distribution of the human proteome. Extracellular proteins were slightly underrepresented, as anticipated based on our experimental design focusing on intra-cellular proteins. Overall, the dataset quantified a substantial proportion of the canonical human proteome (∼80%), while also providing high protein sequence coverage with a median of 50% (**Figure 1E**). This implies that more than half of the protein amino acid sequence was detected in half of the quantified proteins. Comparison of coverage across functional protein families (**Figure 1F**) revealed that more than 90% of protein kinases ^25^, protein phosphatases, transcription factors (TFs), oncogenes, and tumor suppressors^26,27^ were quantified with >50% sequence coverage. However, TFs showed slightly lower sequence coverage of 35%, indicating that TFs are generally lower in abundance compared to other proteins. Importantly, nearly the entire human kinome was covered with ∼500 protein kinases identified in total, without any observable bias in protein detectability or sequence coverage across kinase families (**Figure 1G**).

#### Deep Phosphopeptide library reveals cell line specific signaling

From the nDIA analysis of the phosphopeptide-enriched samples, we generated a very deep phosphoproteome, with a mean of 30,498 class I phosphosites identified per file (from 22,740 to 39,490, **Suppl. Fig. 1B**). Altogether, the resulting phosphopeptide spectral library contains over 500,000 phosphopeptide precursors spanning 13,891 protein groups and representing 223,887 distinct phosphosites (**Figure 2A, Suppl. Fig. 3**). The majority of sites were detected on singly-phosphorylated peptides (multiplicity of one) and were localized with high confidence based on the DIA-specific PTM site localization score algorithm ^20^, with almost two-thirds assigned a site localization probability ≥ 0.99. Overall, the detected phosphosites consisted of 66% phosphoserine (pS), 25% phosphothreonine (pT), and 9% phosphotyrosine (pY) (**Figure 2B**). While the proportion of phosphotyrosine is high in the combined library compared to the expected fraction of a few percent ^18^, individual cell lines displayed an average of 75% pS, 20% pT, and 5% pY, which after filtering for valid values shifted to 81.5% pS, 15% pT, and 3.5% pY. (**Figure 2C**). Interestingly, we find that pY and pT proportions are consistently higher at the expense of pS in both filtered and unfiltered data, indicating that non-overlapping cell type-specific phosphosites increase pY and pT representation in the merged dataset.

**Figure 2:**
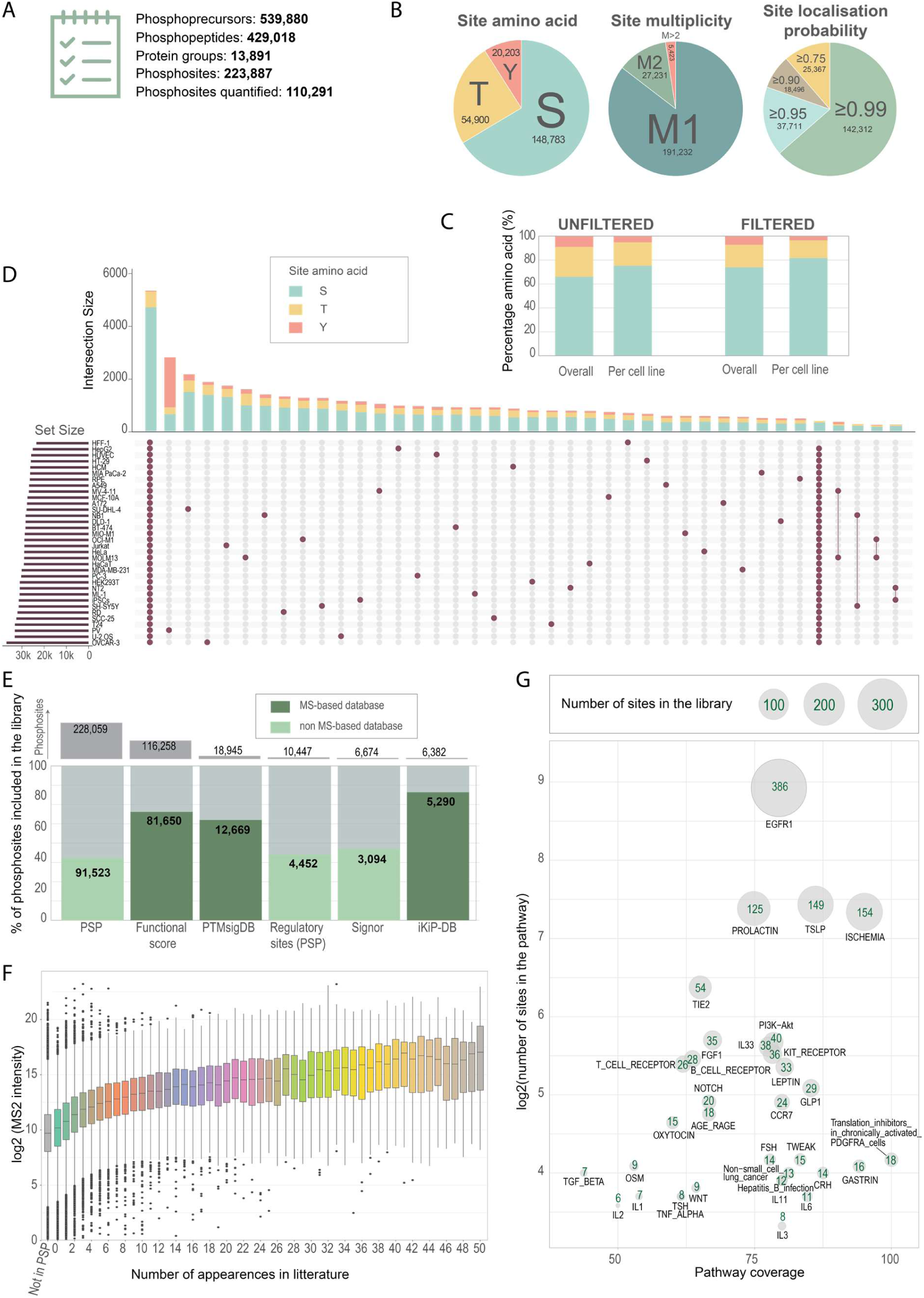
The deep phosphoproteomics library shows high coverage on public databases. A-Overview of the depth of the resulting phosphopeptide spectral library. B- Site amino acid, site multiplicity, and site localization probability distributions over the generated data set. C- Bar plot showing the different proportions of pS, pT and pY between the overall dataset and in each cell line individually, in both the unfiltered (ie. library) and filtered (ie. after valid value filtering) datasets. D- Upset plot presenting the phosphosite overlap across all 33 human cell lines analyzed and the pervanadate treatment. The color of the bars represents the relative proportion of each phosphorylated amino acid. E- Barplot representing the proportion of phosphosites overlapping between our empirical phosphopeptide spectral library and public databases. Darker bars represent the overlap with databases generated using mass spectrometry. The top grey bars represent the total number of phosphosites in the database. F- Relation between the measured MS2 intensity of phosphosites and their number of appearance in MS-based literature according to PSP. G- Coverage over curated signaling pathways from PTMSigDB ^31^. The y axis represents the total size of the pathway and the x axis displays the phosphosite coverage over the pathway with the library. The number of site overlapping between the library and the pathway are shown in green.

The largest intersection across cell lines corresponds to phosphosites shared across all cell lines, reflecting a core signaling set of phosphosites with very low pY content (1%) (**Figure 2D**). In contrast, the majority of additional intersections were unique to individual cell lines rather than shared among subsets. The largest unique intersection corresponded to the pervanadate-treated DLD-1, where over two-thirds of those unique sites corresponded to pY, confirming the effectiveness of the treatment. The proportion of pY in the further one-cell-line intersections was high, with an average of 11%. Additionally, the unique sites per cell line were biased toward the lower intensity range compared to shared sites (**Figure 2D, Suppl. Fig. 4**), which can be consistent with an increased detection of low-abundant phosphosites, including phosphotyrosines. While this could reflect genuine cell line–specific signaling where tyrosine phosphosignalling is used for cell line-specific functionality, it cannot be excluded that it could be an artifact related to FDR control across searches performed separately and subsequently merged. Taken together, the dataset captures a broad landscape of conserved and cell-line-specific signaling.

#### The dataset shows high phosphosite coverage on public databases

To further benchmark the depth of our dataset and assess the quality of our phosphosite detection, we overlapped our dataset with publicly available phosphoproteomics databases. We included (i) PhosphoSite Plus (PSP) ^29^, the largest PTM database with > 200,000 phosphosites, (ii) the manually-curated regulatory sites available in PSP, (iii) Signor ^30^, a database containing experimentally validated causal interactions including specific phosphosites involved in activation or inhibition of a protein, (iv) PTMsigDB, a site-specific curated MS-based data set ^31^, (v) the functional score data set, generated from MS-based available data ^4^, and (vi) the iKiP-DB regrouping in vitro kinase reaction including kinase substrate relationships ^32^. We found that over 60% of the detected sites in our full phosphoproteomics dataset overlapped with the MS-based databases, whereas it decreased to ∼40% overlap for non-MS-derived resources (**Figure 2E**). In total, 81,650 phosphosites were shared with the functional score dataset and 91,523 with PSP, indicating that a large fraction of the detected sites are represented in existing databases. Among the sites annotated in PSP but not detected in our dataset, over half lacked MS-based evidence in the literature and showed an enrichment for phosphotyrosine (**Suppl. Fig. 5**). Many of these sites represent unpublished, in-house generated phosphotyrosine-specific phosphoproteomics datasets based on low-resolution ion trap MS/MS data that may correspond to very low-intensity events ^33,34^, sites that are intrinsically difficult to detect and reproduce by MS, or potential artifacts. Conversely, the fraction without prior MS evidence drops substantially when focusing on the detected sites in our dataset which are overlapping with PSP (**Suppl. Fig. 5**). As site intensity correlates with the number of literature reports in PSP (**Figure 2F**), deep and sensitive phosphoproteomics enabled the detection of low-intensity sites with no previous literature evidence.

We next assessed coverage of phosphoprotein signaling pathways using curated PTMsigDB signatures (**Figure 2G**). Almost all pathways displayed ≥ 50% coverage, meaning that over half of their curated phosphosites were detectable in our spectral library. We found that coverage generally correlated with pathway size, reaching particularly high values for extensively studied signaling pathways such as EGFR, where 386 sites were quantified, while only 7 sites were covered for the TGF-β pathway, with a pathway coverage slightly below 50%. As both EGF and TGF-β ligands are present in serum^35^, and their respective receptors are expressed in the proteome, the serum stimulation should have made it possible to cover both pathways, however, curated pathway databases remain incomplete, as illustrated by the limited TGF-β pathway annotation. At the gene level, both pathways displayed similar levels of phosphosite coverage, including the presence of known regulatory sites (**Suppl. Fig. 6**). Altogether, the inclusion of diverse cell lines with distinct signaling programs contributed to broad pathway representation.

### II. Deep experimental spectral library increases the depth of phosphoproteomes

To evaluate the utility of our large-scale empirical spectral phosphopeptide library, we benchmarked its performance on independent samples and across different chromatographic gradients. We generated proteome and phosphoproteome datasets from a heterogeneous pool of cell types derived from an uncontrolled differentiation of induced-pluripotent stem cells (iPSCs), which was not represented in the library. The differentiated cells were pooled, proteins were extracted and digested, and resulting peptide mixtures were enriched for phosphopeptides. Six technical replicates were analyzed by nDIA across four chromatographic conditions: high-flow gradients of 30 SPD (matching the library conditions) and 100 SPD, as well as low-flow whisper gradients of 80 SPD and 120 SPD (**Figure 3A**). All datasets were searched in both Spectronaut and DIA-NN, comparing empirical spectral library-based searches with the corresponding library-free approaches.

**Figure 3:**
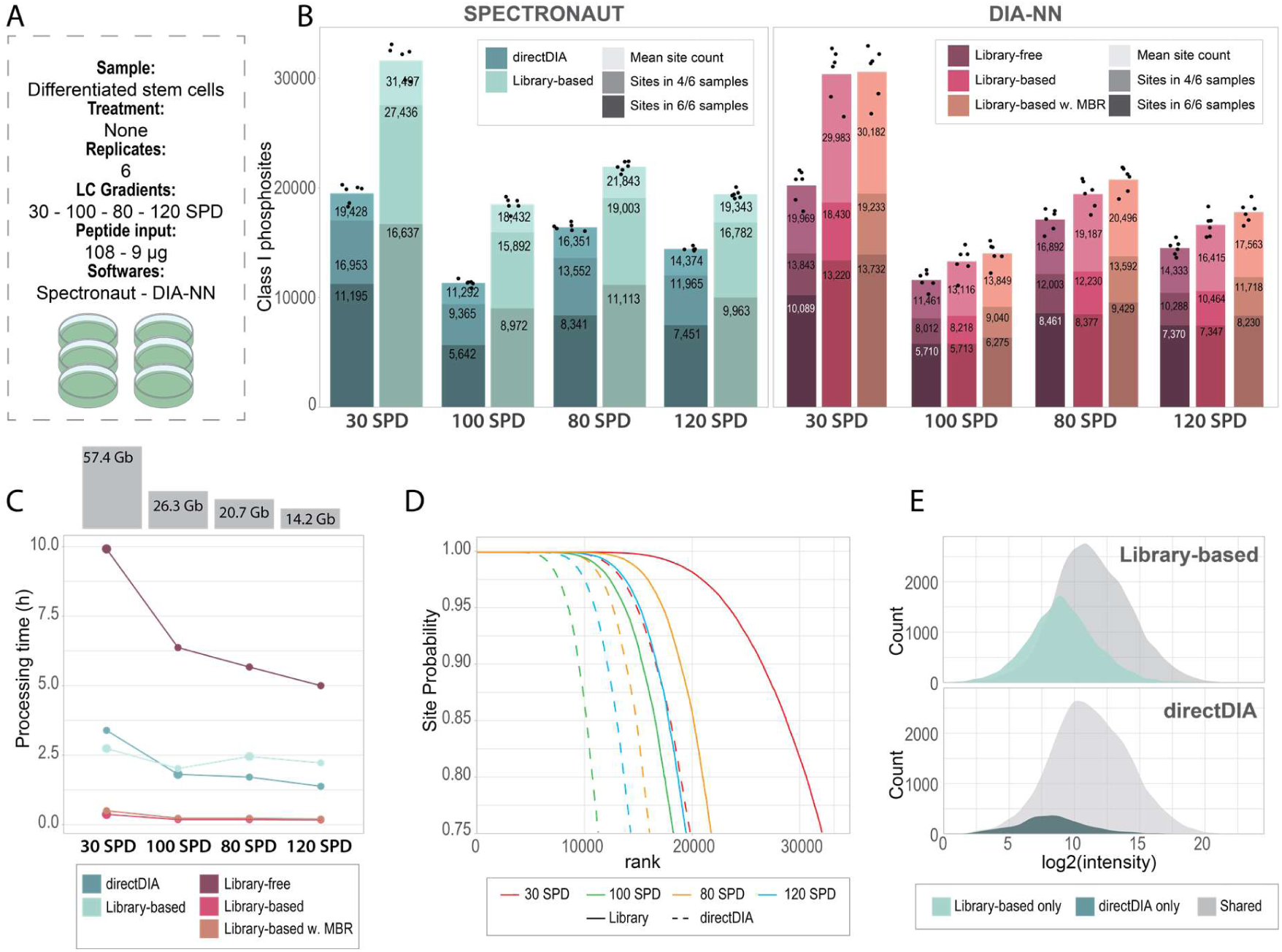
The library-based approach increases phosphosite coverage while reducing computational search time. A- Overview of the experimental design where differentiated stem cells were analyzed on different LC gradients. B- Class 1 phosphosite coverage on four different gradients (30, 60, 80, and 120 SPD). The dots represent the site count per replicate. The lightest bar displays the mean site count across the 6 replicates, the second bar displays the site count after filtering for 4/6 valid values and the darkest bar shows the site count with full completeness across the replicates. SPD - samples per day. C- Line plot displaying the processing time of the searches carried out. The grey bars represent the total file size of the 6 replicates searched. D- Rank plot of the quantified phosphosites based on their localization probabilities using Spectronaut. Each color stands for a different LC gradient, while the full lines represent search outputs using the phosphopeptide spectral library, and the dashed line displays the search results from the directDIA approach. E- Intensity distribution plots using the 30 SPD experiment searched with Spectronaut.

We found that the use of the empirical phosphopeptide library increased phosphoproteome depth compared to library-free approaches regardless of gradient length and search engines, both before and after filtering of the dataset for completeness (**Figure 3B**). In Spectronaut, the benefit of our deep experimental spectral library was particularly pronounced, with a 60% increase in quantified phosphosites under high-flow LC conditions and a 40% increase under low-flow LC conditions when requiring ≥ 4/6 valid values. In DIA-NN, the empirical library mainly improved the median number of quantified phosphosites, whereas the gain after filtering for completeness was minor (1-2.5%). A stronger effect (∼ 30% increase) was observed for samples acquired with the 30 SPD gradient. The close agreement between DIA-NN empirical and in silico searches likely reflects the similarity of the underlying search strategies, and the high quality of the spectral predictions for phosphoproteomics displaying high similarity with empirical spectra (**Suppl. Fig. 7**).

The largest gains from the empirical library were obtained when the LC gradients matched those used for library generation, likely due to improved retention time alignment (**Suppl. Fig**. **8**). Nevertheless, when applied across different LC gradients, retention time correlations remained high (> 0.99), supporting the broader applicability of the library. Overall, Spectronaut provided more phosphosites quantified compared to DIA-NN, which can be attributed to inherently different search strategies and site localization algorithms, but also to different FDR control strategies and cutoff metrics.

We compared computational search times for both library-based and library-free approaches in Spectronaut and DIA-NN (**Figure 3C**). In DIA-NN, the empirical library caused a drastic reduction in search time, which correlated with dataset size. For example, processing time for the six 30 SPD files decreased from 9 h 55 min to just 22 min. Across all gradients, search time was reduced by ∼30-fold on average, reflecting the reduction in spectral library size from 46,686,794 precursors in the in-silico-generated phosphopeptide library to 539,880 in the empirical spectral library. In Spectronaut, however, the library use had a minor effect on search time, even slightly increasing for small datasets. Nevertheless, as dataset size increased, the trend reversed and the library use became beneficial. We expect that this benefit would be more pronounced with larger datasets, where more files would also enable the generation of a larger, more comprehensive Pulsar spectral library, likely reducing the difference in depth between library-based and directDIA. Overall, these data demonstrate that the generated empirical phosphopeptide library improves both quantification depth and computational efficiency across search strategies.

Furthermore, we observed that the increased depth afforded by the empirical spectral library mainly corresponded to phosphosites with high site localization scores (**Figure 3D**). For example, in 30 SPD directDIA, the number of phosphosites quantified with a site localization score of 0.75 matched the number of sites quantified at a cutoff of 0.98 using the library, which contains high-quality spectra with well-localized phosphosites. This suggests that a key limitation of the directDIA approach is the difficulty in confidently localizing PTM sites, whereas the library enables more precise localization. Furthermore, unique sites quantified using the empirical library tend to be of low abundance (**Figure 3E**). Therefore, we can hypothesize that library-based searches could enhance phosphosite identification and localization in experiments with limited input amounts.

### III. The library provides increased depth in sensitive phosphoproteomics experiments

Next, we assessed the potential of the empirical phosphopeptide spectral library to improve coverage of very low-input phosphoproteomics experiments. To test this, we analyzed EGF-stimulated HeLa cells using peptide input amounts ranging from 1 to 20 μg followed by Zr-IMAC-enrichment, and analyzed the resulting phosphopeptide mixtures by nDIA with the 80 SPD and 120 SPD whisper-zoom LC gradients to maximize sensitivity ^36^ (**Figure 4A**). Across all input amounts, library-based searches yielded deeper phosphoproteome coverage, both in terms of the mean number of phosphosites per condition and the number of phosphosites consistently quantified, with an average increase of 18% across all conditions (**Figure 4B, Suppl. Fig. 9A**). Notably, coverage saturated at 10 μg peptides on the 80 SPD gradient, with consistent quantification of over 20,000 phosphosites.

**Figure 4:**
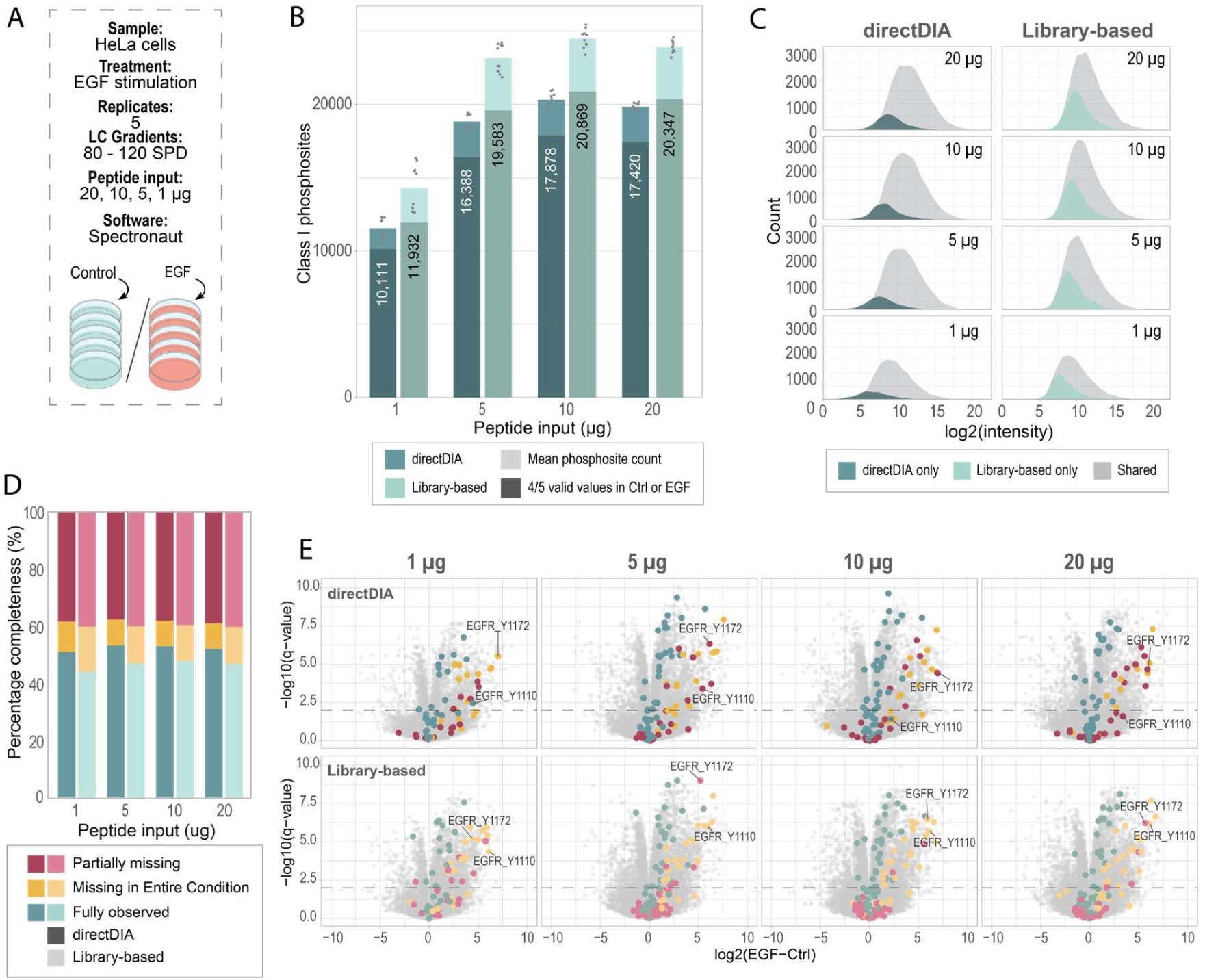
Using a comprehensive phosphopeptide spectral library improves the depth in sensitive phosphoproteomics. A- Overview of the experimental design of the EGF-stimulation scale down on HeLa cells. B- Class 1 phosphosite coverage on peptide input amount ranging from 1 to 20 μg. The dots represent the site count per replicate. The lightest bar displays the mean site count across the 5 replicates in the control and EGF stimulated conditions (N=10), and the darkest bar shows the site count after valid value filtering for ⅘ valid values in the control or stimulated experiments. The number displayed corresponds to the site count after filtering. C- Intensity distribution through the scale down. D- Distribution of the type of data missingness during the scale down in directDIA and using the spectral library. Fully observed values are phosphosites with valid values in all replicates in the control and EGF-stimulated experiments (N=10). Missing in an entire condition are defined by 0 or 1 valid value in the control or the EGF-stimulated experiment, and 4 or 5 valid values in the other condition. All other phosphosites are annotated as partially missing. E- Volcano plots across the scale down for both search approaches tested. Color points represent phosphosites of the canonical EGFR pathway. Each color represents the type of missingness assigned to the site in the given experiment (see legend in D). Horizontal dotted black lines represent a cutoff at 1% FDR.

In the scale-down experiment, unique phosphosites in both library-based and library-free searches were generally of low abundance, with a greater number of unique sites detected using the library due to its higher depth (**Figure 4C, Suppl. Fig. 9A**). In directDIA, the intensities of unique phosphosites followed a normal distribution extending beyond the range of shared values. Quantification was largely consistent between approaches, with a Spearman correlation of ∼0.85 (**Suppl. Fig**. **9B**), although differences in the fragments used for quantification may explain minor variation.

Investigating the missingness across both methods revealed that directDIA generally provides higher data completeness compared to library-based searches (**Figure 4D**). The spectral library-based approach detected more sites considered missing in a full condition, in line with the experimental design, which consisted of EGF-stimulated cells. After left-censored imputation, t-tests were applied to each condition, and both approaches successfully identified canonical EGFR activation sites (**Figure 4E, Suppl. Fig. 9C**). Notably, library-based searches provided more consistent detection of these activation sites at lower input amounts, while directDIA detected some sites partially at low-input. For example, the EGFR autophosphorylation site Y1110 was either fully or partially observed in directDIA but was detected as missing across the entire control condition and fully present in the stimulated condition using the library-based approach. This suggests that higher completeness in directDIA may reflect noisier integration or differences in cutoff strategies in low-input phosphoproteomics, with overall lower scored precursors leading to lower score cutoffs. Regardless, quantification was not affected by the search strategy used and showed robust fold changes. We observed strong fold-change correlations between both library-based and library-free methods, with slightly lower correlation for phosphosites labelled as partially missing sites (**Suppl. Fig. 9B**).

Overall, this analysis showed that empirical spectral libraries enhance the depth and site localization confidence in low-input phosphoproteomics experiments. When generation of sample-specific deep libraries is not feasible, for example due to low amounts of starting material available, our large-scale empirical library represents a reliable alternative for comprehensive and fast phosphoproteomics profiling.

### IV. Site stoichiometry calculation draws a clear picture of early signaling events upon serum stimulation

Determining not only the relative quantification but also the occupancy or fractional stoichiometry of a phosphosite is highly informative, as a large stoichiometry for a dynamically regulated site strongly suggests that it is functionally important ^37^. We therefore aimed at calculating phosphosite stoichiometry to provide an absolute measure of the phosphorylation occupancy^19,38–41^. Stoichiometry represents the fraction of a site that is phosphorylated and the calculation requires high depth and data completeness, with both the phosphorylated and non-phosphorylated forms of a peptide detected in control and stimulated conditions. Additionally, phosphorylation can only be calculated when a change is observed between both conditions.

Stoichiometry was calculated in each of the 31 cell lines, excluding SCC-25 and PC-3, which sowed mycoplasma contamination, reflecting the effect of serum stimulation^19^. On average, stoichiometry could be determined for 1,580 phosphosites on average per cell line spanning a wide range from 474 to 2970 sites, reflecting variability in cell stimulation efficiency and responsiveness. In total, stoichiometry was determined for 19,378 sites encompassing 15,443 pS (80%), 3,109 pT (16%) and 826 pY (4%) (**Figure 5A**). The stoichiometry for majority of sites were cell-line specific (**Figure 5B**), likely due to missingness arising from the requirement to match phosphoproteome and proteome data at the modified peptide level, where most stoichiometry-quantified sites corresponded to phosphopeptide with high MS2 intensities (**Figure 5C**). Across all cell lines, stoichiometry and phosphopeptide intensity showed only a weak positive correlation, suggesting that the two metrics capture distinct aspects of the data (**Figure 5D**).

**Figure 5:**
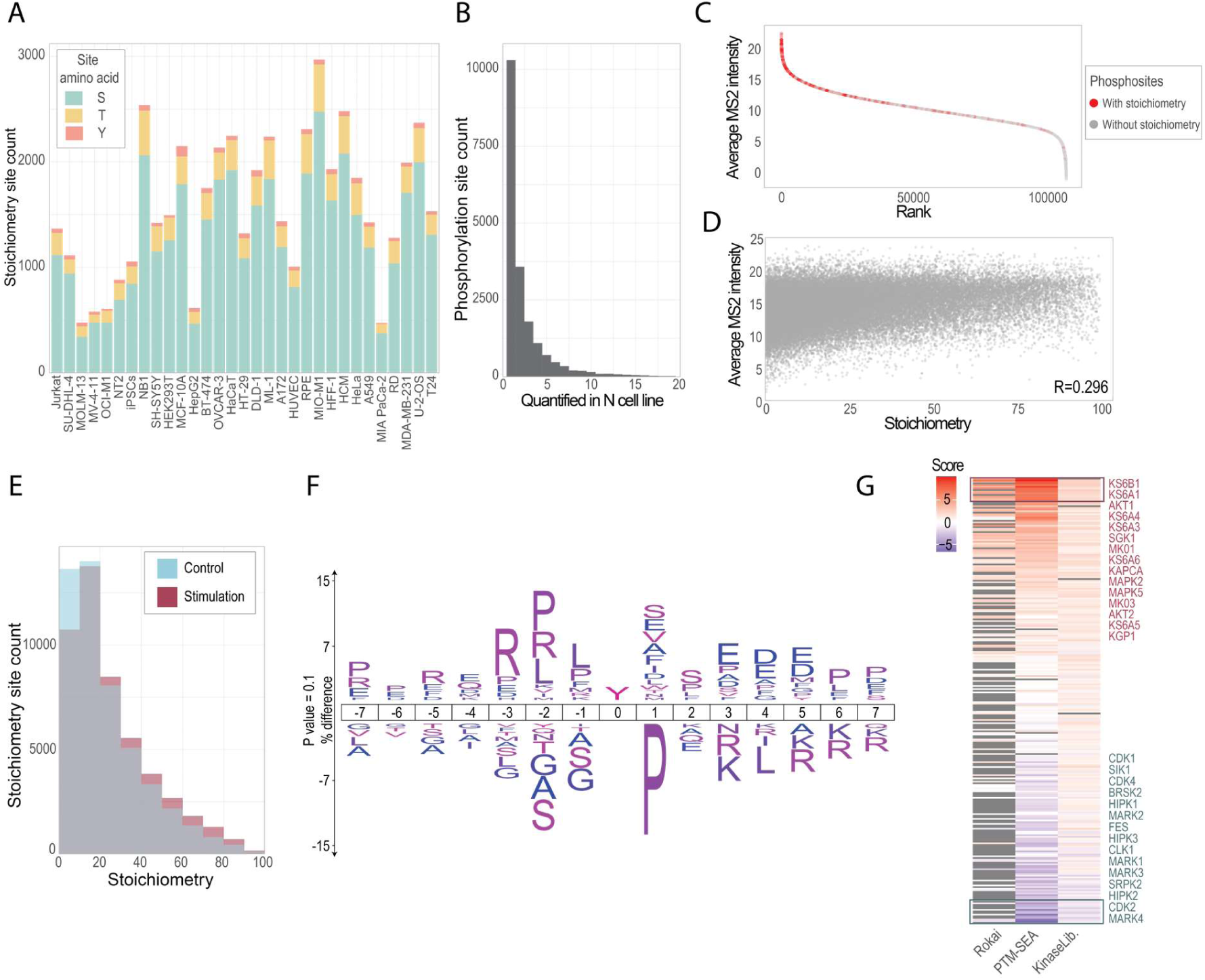
Stoichiometry calculation allows for a clear overview of early signaling events upon serum stimulation. A-Overview of the stoichiometry site count per cell line in the serum stimulated condition relative to the control. B- Histogram representing the overlap between calculated sites across cell lines, where the x axis represents the number of cell lines across which a given site had its stoichiometry calculated. C- Rank plot of the measured average MS2 intensity across all cell lines per phosphosite. Sites where stoichiometry could be calculated are displayed in red. D- Correlation between the average MS2 intensity per cell line and the corresponding calculated site stoichiometry for the given cell line with Spearman correlation coefficient displayed on the graph. E- Distribution of the computed stoichiometry values in the control and serum stimulated conditions. F- IceLogo of the phosphorylated sites (top plot) compared to dephosphorylated sites (bottom plot) upon serum stimulation ^45^. G- Heat map of the kinase activity analysis computed using Rokai, PTM-SEA using the modified iKiP-DB and the Kinase Library. Scores were calculated based on the difference between the stoichiometry in the serum stimulated condition and the control. Kinases represented in red represent the top 15 most activated kinases and the ones in blue represent the top 15 most inactivated kinases.

Stoichiometry values were generally low (< 20% occupancy) and increased upon serum stimulation, with distributions shifted toward higher occupancy values (**Figure 5E**), reflecting the activation of signaling pathways. It was then possible to pinpoint exactly the phosphosites being phosphorylated or dephosphorylated upon serum stimulation. Phosphosites with a positive fold change (phosphorylated) were enriched in basophilic sequence motifs with an overrepresentation of arginine in positions -3 and -2 to the phosphosite, which is the characteristic for substrates of kinases from the AGC family, including PKA, PKB or S6K (**Figure 5F**). A moderate enrichment in tyrosine phosphorylation was also observed. In contrast, the dephosphorylation motifs were enriched in proline directed phosphosites with proline at position +1, typically targeted by MAPKs and CDKs, indicating phosphatase-mediated deactivation of these pathways. To further resolve kinase activity changes, kinase–substrate enrichment analysis was performed on stoichiometry fold changes between stimulation and control (**Figure 5G**). The top 15 most serum-activated kinases identified by three independent kinase inference algorithms included ribosomal protein kinases (KS6B1, KS6A1, KS6A4, KS6A3, KS6A6, KS6A5), PKB (AKT1, AKT2) and MAPK (MAPK1 (MK01), MAPK2, MAPK3 (MK03), MAPK5). Conversely, the most strongly deactivated kinases implicated by substrate dephosphorylation were MARK family members responsible for the phosphorylation of microtubule associated proteins, promoting microtubule dynamics^42–44^. Moreover, cell cycle regulating CDKs (CDK1, CDK2 and CDK4) were also among the most inactivated kinases.

Altogether, our data indicate that a 10-minute serum-stimulation induces activation of mitogenic and growth-related signals, activating protein synthesis and pro-survival signaling, while cell cycle progression promoters are inactivated. Thus, the phosphoproteome stoichiometry changes observed by short-term serum stimulation revealed metabolically and translationally active but not proliferative cellular state, which will likely shift at a later time point. While the immediate early signaling response suggests cell growth and protein production, later signaling would be focused on promoting re-entry into the cell cycle.

### V. Deep proteomics profiles enable functional clustering of the cell lines

We next explored functional grouping of the analyzed human cell line proteomes. Principal component analysis (PCA) of the log2-transformed protein intensities revealed a clear separation between suspension (i.e. blood cancer) and adherent cell lines (**Figure 6A**). Additionally, non-cancerous and cancerous cells showed partial separation along the first principal component (PC1), with a distinct cluster likely corresponding to mesenchymal-like cells. To further refine the classification, we applied consensus clustering to the proteome data, identifying six clusters based on 14,570 quantified protein groups (**Figure 6B**). Functional annotation of each cluster was performed using String ^46^ by analyzing the top 100 most differentially expressed proteins per cluster, defined as those with the highest fold change relative to all other clusters, enabling functional characterization of the distinct cell line groups (**Figure 6C, Suppl. Fig. 10**).

**Figure 6:**
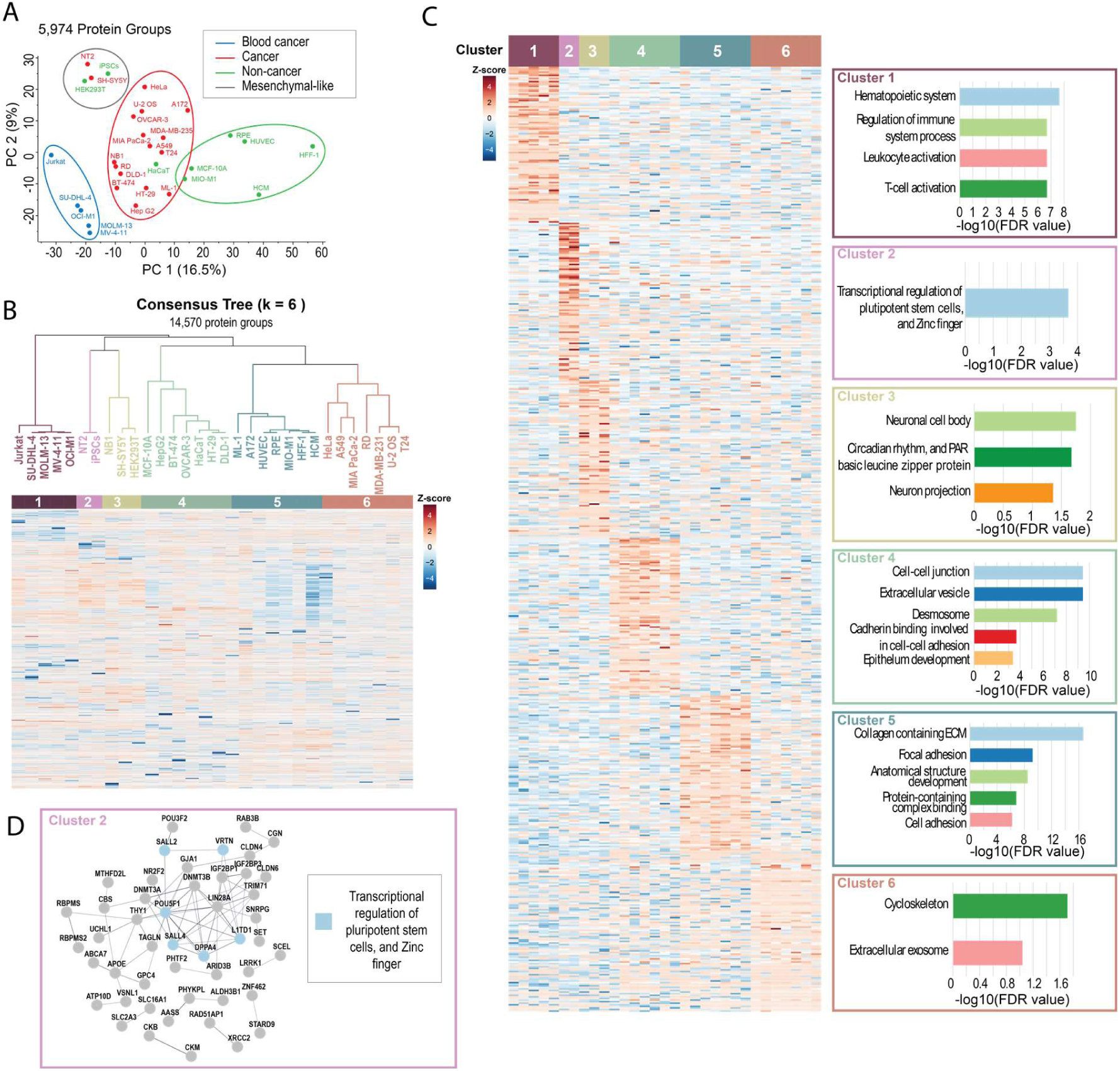
Functional analysis across cell lines shows distinct clusters. A- PCA plot based on the proteome data filtered for full completeness. B- Cell line clustering using consensus clustering on 14,570 quantified protein groups. Missing values were imputed using KNN. C- Heat map with the top 100 protein groups per cluster that were the most differentially overexpressed compared to the background of all protein groups. The results on the gene set enrichment of those proteins were carried out on Cytoscape. D- Network analysis of the top 100 protein groups of cluster 2, showing stem cell-specific proteins.

Functional analysis of the clusters revealed distinct biological characteristics. A first cluster encompassing the five blood cancer cell lines was characterized by the overexpression of proteins associated with the hematopoietic system and immune cell activation. Cluster 2 contained pluripotent stem cell-like cell lines, including the iPSCs and the NT2 pluripotent human embryonal carcinoma cell line, with NT2 exhibiting very similar proteomic features to iPSCs. Both cell lines showed overexpression of a dense network of proteins involved in transcriptional regulation of pluripotency, including OCT4, SALL4, SALL2, DPPA4, and LIN28A ^47,48^ (**Figure 6D**). The third cluster corresponded to neuronal-like cell lines, comprising two neuroblastoma lines (NB1 and SH-SY5Y) and HEK293T, which has previously been shown to resemble neuronal cells despite being of embryonic kidney origin ^49,50^. Differentially expressed proteins were associated with neuronal cell bodies and projections, though enrichment FDR was low, likely because cluster 2 also exhibited neural characteristics, reducing uniqueness. Cluster 5 grouped most non-cancerous cell lines, with the exception of ML-1 (follicular thyroid carcinoma) and A172 (non-tumorigenic glioblastoma). Proteins overexpressed in this cluster formed dense networks related to extracellular matrix (ECM) organization, including terms such as “focal adhesion” and “cell adhesion,” reflecting the capacity of cells to anchor, interact, and build structural networks in their environment.

Finally, two different clusters, cluster 4 and cluster 6, containing epithelial cancerous cell lines (with the exception of MCF-10A and HaCaT) were identified. Cluster 4 showed a high degree of epithelial organisation with a strong enrichment of the terms “cell-cell junction”, “desmosome”, “cadherin binding”, and “epithelium development”. This suggests that those cell lines maintain tight intercellular adhesion, allowing them to form organised and cohesive cellular layers. On the other hand, cluster 6 only showed enrichment of broad terms with weak statistical significance. This likely originates from the cluster being composed of cell lines that are more different from each other compared to the variability observed in other clusters. Therefore, the difference between the both clusters comes from the variable epithelial characteristics of the cancer cell lines, in agreement with their in vitro growing patterns, indicating a potentially less aggressive cancer group with maintained structural organisation, and a likely more invasive cancer cell line group showing a less adherent phenotype.

### VI. Kinase activity analysis and correlation with drug sensitivity uncover therapeutic vulnerabilities

Having established functional groupings and lineage-specific features across the panel of cell lines, we next attempted to infer kinase activity patterns. Inferring kinase activity is essential to translate phosphosite-level changes into functional signaling insights by identifying which kinases drive the observed dynamics ^51–53^. This is particularly relevant in cancer research, where aberrant kinase activities often drive tumor progression and represent promising therapeutic targets ^54^.

Kinase activity analysis typically relies on the detection of phosphosites on downstream substrates using established kinase-substrate relationships or kinase-specific motif patterns ^31,55–61^. However, these approaches present challenges for closely related kinases, particularly tyrosine kinases, that share a large proportion of their downstream substrates ^57,62^. In such cases, computational kinase activity inference analysis tools such as Rokai ^59^, PTM-SEA ^31,32^, or the Kinase Library ^62^ often lack resolution. After computing inferred kinase activity scores on our dataset using these three approaches, we observed limited resolution among tyrosine kinases, leading to false-positive signals of certain kinases (**Figure 7A**). For instance, the well-studied EGFR was expected to be active in epithelial cells only and is not expressed in non-epithelial cells, as confirmed by our proteome data, where EGFR is absent in all of the blood cancer cell lines. Nevertheless, all three inference tools report a kinase activity score for EGFR in the blood cancer cell lines, in some cases even assigning positive activity, implying higher EGFR activity compared to the background of all cell lines.

**Figure 7:**
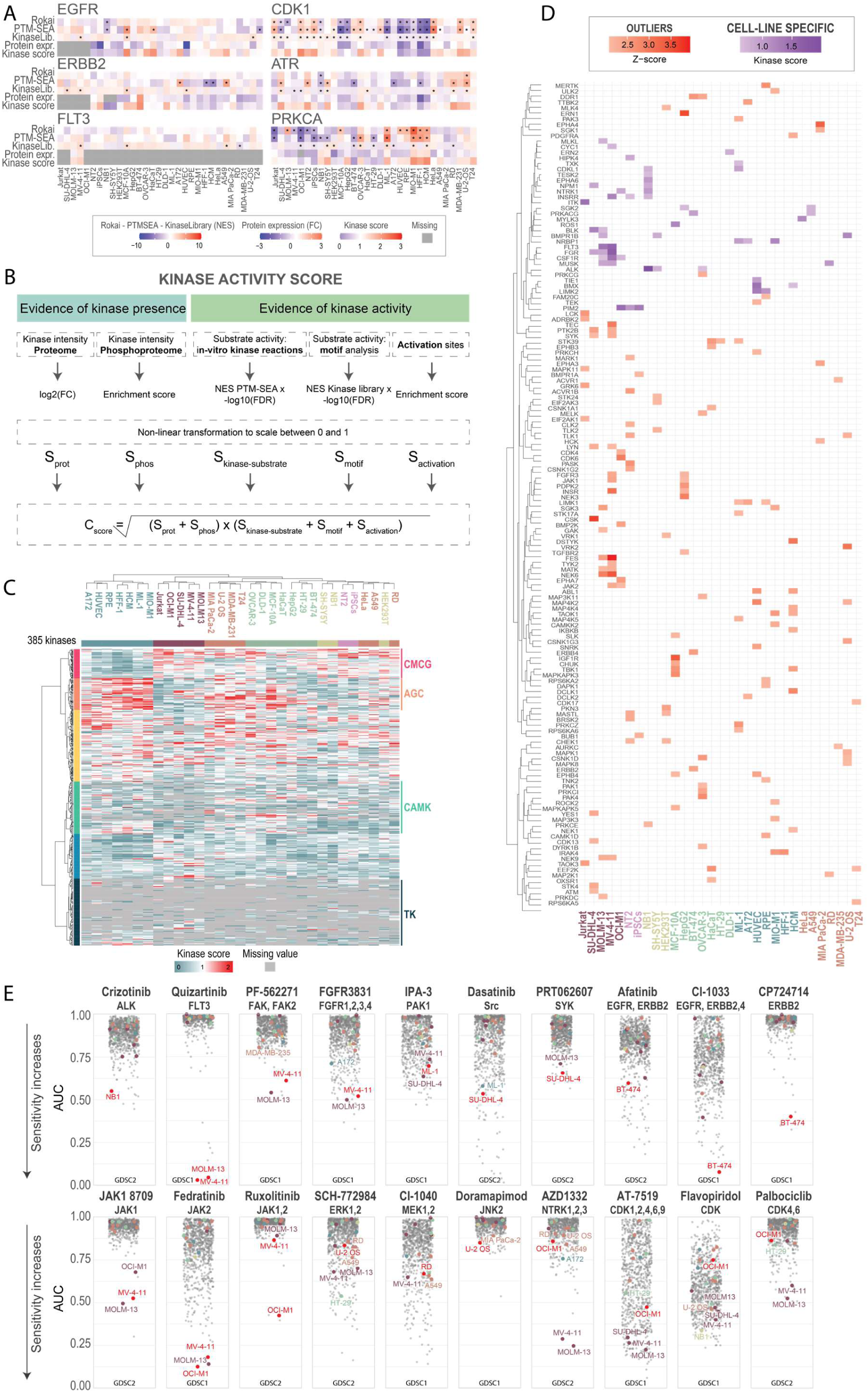
A combined kinase score allows to identify cell-line specific kinase activity landscapes. A- Heatmaps displaying the inferred kinase activity calculated across cell lines with RoKai, PTM-SEA using the iKiP-DB, the Kinase Library, the kinase protein expression and a combined kinase activity score for six kinases, including three tyrosine kinases (on the left). The stars represent statistical significance at 5% FDR. B- Flow chart of the combined kinase activity score. C- Heatmap of the combined kinase score across cell lines. D- Heatmap displaying the kinase-cell line associations. The red squares represent associations established by identifying kinase activity outliers, and the purple squares display associations defined by cell line-specific kinase activity. E- Drug sensitivity screen data from GDSC1 or GDSC2 over different kinase inhibitors. Each red dot represents a kinase-cell line association. Colored dots represent cell lines in the panel on which the kinase inhibitor was tested, and grey dots show other cell lines in the drug test panel. The name of the kinase inhibitor is on the top row, and the targeted kinase(s) are on the bottom row.

Likewise, ERBB2 is known to be expressed in epithelial cells with a high protein abundance level in ERBB2-positive breast cancer cell lines such as BT-474. Yet, all three kinase inference methods computed positive ERBB2 activity scores in blood cancer cell lines, and failed to identify that BT-474 has significantly higher ERBB2 activity compared to the other cell lines. Finally, FLT3, which is specifically expressed in hematopoietic progenitor cells and is constitutively active in MOLM13 and MV-4-11 due to a FLT3 internal tandem duplication (ITD) mutation ^63^, should therefore exhibit higher kinase activity in these cell lines. However, this is not captured using the kinase activity inference tools (**Figure 7A**).

To increase the resolution and fully exploit the depth of our dataset to obtain a comprehensive overview of kinase activities across the cell lines, we developed and applied a combined kinase activity score by incorporating information from the paired pan-proteome and phosphoproteome, capturing kinase activation and substrate phosphorylation (**Figure 7B**). Our combined kinase activity score (Cscore) requires the presence of a given kinase in the proteome and/or phosphoproteome and its activity inferred in the phosphoproteome to compute a score. The kinase activity is derived from three complementary elements. Two of these scores are based on substrate phosphorylation and are directly computed through pre-existing tools: one using established kinase-substrate relationships from in vitro kinase reaction experiments ^64^ complemented with PSP annotations^31,32^(**Suppl. Fig. 11**), and the other based on sequence motif analysis ^62^. The third activity score takes into account the detection of activation sites on the kinase.

Altogether, we were able to compute kinase activity scores for 385 kinases across 31 human cell lines (**Figure 7C**). Our Cscore provides improved resolution for tyrosine kinases, with missing values appropriately assigned to kinases not expressed. Importantly, the Cscore correctly scores the directionality of a kinase activity state in individual cell lines, as demonstrated with the zero activity score of both EGFR and ERBB2 in suspension cells and the high activity score of ERBB2 in BT-474 (**Figure 7A**). For other non-tyrosine kinases, the computed scores were consistent with previously assessed kinase activity using existing tools in both scale and directionality. For example, CDK1 and ATR showed higher activity in cancer cell lines compared to non-cancerous cell lines.

Clustering the kinase activity profiles revealed six groups of kinases with similar activity patterns across the cell lines. The first cluster was dominated by CMGC kinases (**Figure 7C, Suppl Fig. 12**) and showed low activity in non-cancerous and high activity in cancerous cell lines, likely reflecting a basal, non-proliferative signaling state in non-cancerous cells. Another cluster, enriched in AGC kinases, displayed the opposite pattern, with higher activity in non-cancerous cells. CAMK kinases formed a further cluster, characterized by sporadic detection and cell line-specific activity. An additional cluster was enriched in tyrosine kinases, whose activity was highly specific to individual cell lines, reflecting selective expression and signaling. Additionally, we identified specific kinase with high activity in different functional clusters (**Suppl Fig. 12**) with kinase specifically involved in immune regulation (i.e. BTK, SYK, FGR) in the blood cancer cell lines, or PIM2 in the pluripotent cell cluster, which has been associated with cancer therapy resistance ^65^.

We next attempted to identify kinase-specific therapeutic vulnerabilities, defined as key proteins for a celĺs growth and survival, and validate these hypotheses using large-scale drug screening data from the Genomics of Drug Sensitivity in Cancer (GDSC) project ^66,67^. We hypothesized that cell lines with high activity of a given kinase would be more sensitive to inhibitors targeting that kinase ^68^.

Cell-line specific vulnerabilities were investigated through the identification of kinase-cell line pairs in which a kinase in a given cell line either showed activity substantially higher than its activity distribution across all other cell lines or it was considered cell line-specific when active in four or fewer cell lines. Kinases in both categories were treated as potential therapeutic targets, resulting in 202 kinase-cell line associations (**Figure 7D**). We then focused on associations involving cancer cell lines and kinases with available and specific small molecule ATP-mimetic inhibitors, substantially narrowing the set of candidate targets. Depending on data availability, we assessed the sensitivity of these cell lines with corresponding inhibitors using the measured area under the curve (AUC) values from the GDSC dataset.

Several of the identified kinase-cell line associations corresponded to known drug sensitivities (**Figure 7E**). For instance, NB1 (neuroblastoma) with high ALK activity score exhibited high sensitivity to ALK inhibition, and cell lines with active FLT3 (MOLM13 and MV-4-11 cell lines) were sensitive to FLT3 inhibitors. Moreover, MV-4-11 cells responded to FAK and FGFR inhibition, ML-1 cells to PAK1 inhibition, and SU-DHL-4 cells to Src and SYK inhibition. Notably, high activities of ERBB2 and ERBB4 were identified as outliers in BT-474 cells, suggesting targetable vulnerabilities. BT-474 cells were sensitive to Afatinib, an EGFR and ERBB2 inhibitor. When treated with CI-1033, which additionally targets ERBB4, BT-474 cells showed increased sensitivity. In contrast, selective inhibition of ERBB2 reduced overall sensitivity but improved specificity, with only a few cell lines, including BT-474, showing a response. Similarly, JAK inhibitors effectively targeted the intended blood cancer cell lines, and MAPK pathway inhibitors (ERK, MEK, JNK) showed strong effects in U-2 OS and RD cell lines, which exhibited among the lowest AUCs.

Some kinase-drug relationships were less straightforward. For example, OCI-M1 cells were associated with high NTRK1 activity, yet NTRK inhibitors were more effective in other cell lines such as MOLM13, MV-4-11, A172, and A549, and showed limited sensitivity in OCI-M1. CDK4 and CDK6 were predicted vulnerabilities for OCI-M1 cell line, and several inhibitors with varying selectivity were tested. Broad-spectrum CDK inhibitors affected multiple cell lines, whereas highly specific CDK4/6 inhibitors had limited effects, and none-selectively affected OCI-M1 cells viability. This highlights the challenge of interpreting drug screens, particularly when drug selectivity is not well described and off-target effects can interfere with the results ^69^.

Overall, these observations support the hypothesis that highly specific and selective kinase inhibitors can validate the predicted cell-specific kinase vulnerabilities. Additional potential targets were identified, but they lacked experimental validation. For instance, MIA PaCa-2 cell line displayed highly specific activity of EPHA3 and EPHA4 kinases, which are implicated in angiogenesis. Inhibiting ephrin kinases could potentially limit pancreatic cancer cell growth, consistent with reports that high ephrin expression correlates with poorer prognosis ^70^ and that EphA3 inhibition reduces tumor growth and progression in pancreatic cancer ^71^. Similarly, ERN1 was an extreme outlier in HepG2 cell line. Given its role in modulating tumor progression in colorectal cancer ^72^, it could represent a potential therapeutic target in hepatic cancer.

## DISCUSSION

In this study, we generated deep proteome and phosphoproteome profiles across a wide panel of 33 human cell lines representing the major tissue-types in the body. With an average of 9,700 protein groups and ∼30,000 class I phosphosites quantified per cell line, this dataset constitutes a unique community resource, which can be directly used as a high-quality experimental spectral library. Notably, the proteome library displays not only high depth but also remarkably high sequence coverage, with over 16,000 protein groups detected and a median sequence coverage of 50%. Previous large-scale studies reported up to ∼17,000 protein-coding genes using diverse approaches such as complementary proteases or extensive offline peptide fractionation ^73,74^. In contrast, our dataset was generated with tryptic peptides using single-shot LC-MS/MS analysis. It remains unclear whether the remaining gap to cover the whole human proteome, estimated at ∼20,000 canonical proteins, reflect limits in our acquisition depth, experimental design, or genuinely undetectable regions of the human proteome. With more than 200 different cell types in the human body, our dataset will also lack cell-type specific proteins from cells that were not part of our panel. For example, the mammalian genome encodes up to 1,000 odorant receptors, which is a large family of G-protein-coupled receptors (GPCRs) found almost exclusively on the cilial membrane of olfactory sensory neurons in the olfactory epithelium ^75^. Since we did not profile this cell type, GPCRs were not covered in our dataset. Moreover, the searches were performed against the canonical reference proteome, and we anticipate the data set to be able to further reveal cell-line-specific protein isoforms or mutations ^76,77^.

Our phosphoproteomics profiling achieved unprecedented coverage within a single unified experiment, with > 223,000 unique phosphosites detected across all cell lines and >110,291 phosphosites consistently and confidently quantified. Approximately ∼90,000 of these sites overlapped with PhosphoSitePlus, including phosphosites without previous MS evidence. High coverage of curated signaling pathways ^30,31^ revealed the power of this large dataset for studying signaling events. Unlike widely available large-scale OMICs datasets, where phosphoproteomics is rarely included, or only available on cancerous material, our panel includes both cancerous and non-cancerous cell lines, offering new opportunities to compare healthy versus malignant signaling states ^8,10,11^.

While the use of a spectral library for DIA-based phosphoproteomics was the standard approach before the emergence of directDIA ^20^, our results showed that a highly complete empirical phosphopeptide spectral library can still substantially improve DIA-based phosphoproteomics, without the need for fractionation. In both Spectronaut and DIA-NN, the empirical library outperformed the respective library-free approaches. DIA-NN’s library-free workflow relies on a large in silico-generated spectral library from a FASTA file, reduces it after a first-pass search, and then performs feature extraction ^78^. Replacing this large theoretical search space with our empirically derived library dramatically narrows the initial search space, which accounts for the modest but consistent gains in identification depth and the striking ∼ 30-fold reduction in computational processing time. The limited unique identifications provided by the library-fee approach further underscore the high degree of completeness of the empirical library. Spectronaut’s directDIA, by contrast, builds a library directly from the DIA data through an initial DDA-like search, and its depth therefore depends strongly on the quality and number of input files available for confident identification and localization of phosphopeptides ^20,79^. In this context, applying our empirical library resulted in a substantial increase in identifications but did not reduce processing time. Given the directDIA workflow, we anticipate that benchmarking on a larger dataset would have narrowed the identification gap and more clearly showed the processing-time benefits of using an empirical library.

Importantly, the empirical library improved performance in high-sensitivity phosphoproteomics experiments with low-input amounts, allowing for an average gain of 18% phosphosites. We observed that directDIA produced an artificially high proportion of complete data, including quantifications of sites that were entirely absent from the unstimulated condition but quantified in all the EGF-stimulated conditions when searched using our experimental library, such as phosphosites on kinase activation loops. This suggests that the primary bottleneck in scaled-down phosphoproteomics is obtaining MS/MS spectra of sufficient quality to confidently identify and localize phosphopeptides. As input amount decreases, spectral quality degrades and statistical cutoffs may become suboptimal ^80^. Using the empirical library alleviates these issues by supplying high-quality reference spectra, thereby reducing the reliance on low-quality MS/MS. Recent advances in ultra-low-input sample-preparation workflows have pushed the sensitivity of phosphoproteomics to unprecedented levels ^15–17^. Integrating these advances with a comprehensive empirical spectral library is likely to further expand coverage and move a step closer to obtaining extensive phosphoproteomes from reduced cell populations.

Determining phosphorylation site stoichiometry adds a crucial quantitative dimension to large-scale phosphoproteomics data. This information reveals not just where phosphorylation occurs or if it is regulated in a given condition, but how much of a protein is modified at each site, which is vital for understanding the functional impact of the phosphorylation on protein activity, signaling pathway regulation, and disease processes. Stoichiometry data enable researchers to distinguish between functionally relevant and incidental phosphorylation events, prioritize therapeutic targets, and build more accurate models of cellular signaling networks. We estimated the stoichiometry for the dynamically changing sites upon ten minutes of serum stimulation, which allowed for the calculation of the fractional site occupancy for more than 19,000 phosphosites in total with 1,580 sites on average per cell line, with variability reflecting differences in stimulation efficiency and cell line-specific responses. Motif and kinase enrichment analyses pointed to strong activation of AGC kinases and MAPKs, whereas CDKs and MARK kinases were predominantly dephosphorylated, suggesting an early signalling response to serum stimulation focused on proliferation rather than cell-cycle progression. Together, these results show that stoichiometry can pinpoint which phosphorylation events are newly gained or lost upon stimulation, providing resolution beyond relative abundance-based phosphoproteomics.

Although stoichiometry offers a valuable complementary perspective on phosphorylation dynamics and has become more feasible with increasing data depth and completeness, the approach remains limited by data sparsity and measurement precision. Each stoichiometry value accumulates the error of four independent intensity measurements (phosphorylated and non-phosphorylated forms in control and stimulated samples), reducing sensitivity for identifying small occupancy changes. Consequently, low-occupancy sites that double upon stimulation may remain within measurement uncertainty, while mid-occupancy sites appear dynamic and lead to an occupancy calculation. Since most sites exhibited low occupancy, unbiased and precise stoichiometry calculation remains a challenge. Focusing on stoichiometry at a smaller scale and with emphasis put on precision, through targeted approaches for instance ^41^, could be helpful for unveiling specific biological insights.

Beyond depth, our dataset also revealed clear cell line-specific signaling states. Functional clustering of the phosphoproteomes showed that signaling programs largely followed cell morphology and lineage. To systematically compare kinase activities, we developed a combined kinase activity score, the Cscore, optimized to discriminate among kinases with shared substrates. This enabled the profiling of 385 kinases across 31 cell lines, revealing clear differences between cancerous and non-cancerous states and highlighting cell line-specific abnormally active kinases. Importantly, because the Cscore’s integrates multiple independent lines of evidence for both kinase presence and activity, the absence of a Cscore can be interpreted as a lack of activity in that cell line. We further correlated these kinase-activity profiles with publicly available drug-sensitivity data and validated cell-line kinase vulnerabilities in several cell lines by their kinase inhibitor responses. Although the validation is limited since many kinase-cell line combinations are not druggable, lack a selective inhibitor, or are absent from the GDSC dataset ^67^, we demonstrated sensitivity on several well-established and undemonstrated associations, suggesting that further vulnerabilities could be demonstrated with drug screens. The limited specificity of kinase inhibitors complicated direct validation, as off-target effects are common and often uncharacterized. Although specificity profiles have been reported for many kinase inhibitors^69^ , they rarely overlap with available cell viability datasets. Nonetheless, this resource can support rational drug testing strategies and combination-therapy design by enabling prioritization of the most active kinases in each cell line.

Overall, we provide a versatile resource that can serve as a high-quality experimental spectral library, support data exploration and mining, guide targeted MS assay development, inform drug-sensitivity studies, and help identify new therapeutic vulnerabilities.

## Material and methods

### 1. Cell culture

All cell lines were cultured at 37°C in a humidified incubator with 5% CO2, and all media were supplemented with 10% fetal bovine serum and 100 μg/mL penicillin/streptomycin (Invitrogen), unless specified otherwise.

A172 (CRL-1620), A549 (CCL-185), DLD-1 (CCL-221), HaCaT (CVCL_0038), HEK293T (CRL-3216), HeLa (CRL-2), Hep G2 (HB-8065), HFF-1 (SCRC-1041), HT-29 (HTB-38), MDA-MB-231 (HTB-26), MIA PaCa-2 (CRL-1420), MIO-M1 (CVCL_0433), ML-1 (ACC 464), NT2 (CRL-1973), RD (CCL-136), RPE (CRL-4002), SH-SY5Y (CRL-2266), T24 (HTB-4), and U-2 OS (HTB-96) were cultured in DMEM (Gibco). BT-474 (HTB-20), Jurkat (TIB-152), MOLM-13 (ACC 554), NB1 (CVCL_1440), and OVCAR-3 (HTB-161) were cultured in RPMI (Gibco). SCC-25 (CRL-1628) and PC3 (CRL-1435) were cultured in DMEM-F12 (Gibco). MV-4-11 (CRL-9591) and OCI-M1 (ACC 529) were cultured in IMDM (Gibco). SU-DHL-4 (CRL-2957) was cultured in RPMI with ATCC modification (Gibco). MCF-10A (CRL-10317) were cultured in DMEM/F12 with glutaMAX (Gibco) supplemented with 5% horse serum (Gibco), 1% (wt/vol) non-essential amino acids (Gibco), 0.01 mg/ml insulin (Sigma-Aldrich), 100 ng/ml cholera toxin (Sigma-Aldrich), 500 ng/ml hydrocortison (Sigma-Aldrich), and 20 ng/ml EGF (Peprotech). HUVEC were cultured in Endothelial Cell Growth Media (EGM-2) (Lonza) supplemented as provided by the manufacturer. HCM (C-12810) were cultured in Myocyte growth media (PromoCell) supplemented as recommended by the manufacturer. Human iPSCs hi12^81^ were expended on laminin521-coated dishes in E8 medium (Gibco) supplemented with 0.1% bovin serum albumin (BSA).

iPSCs differentiation into embryoid bodies (EBs) was carried out by dissociated hi12 cells using TrypLE Express for 5 minutes. TrypLE was removed, and the cells were rinsed once with fresh E8 medium supplemented with 0.1% BSA, scraped, resuspended in fresh medium with 10 μM of Y-27632 (ROCK inhibitor), and cultured in suspension in low-adhesion flasks. After a week, EBs were plated on gelatin-coated dishes in DMEM medium supplemented with 20% (vol/vol) fetal bovine serum, 2 mM l-glutamine and 1% (wt/vol) nonessential amino acids. The medium was replaced with fresh medium every three days.

### 2. Cell lysis and quantification

Each cell line was cultured until 80% confluence in p10 cell culture dishes in 8 replicates. After 48h of culture without media change, the cells were serum-starved for 4 hours. Half of the dishes were serum-stimulated for 10 minutes with the addition of 10% FBS (Fetal Bovine Serum), while the remaining dishes remained unstimulated. Starvation of MOLM13, MV-4-11 and OCI-M1 was performed for 2 hours in medium containing 0.01% BSA. SU-DHL-4 cells were not starved and stimulation was carried out with the addition of and additional 10% of FBS. iPSCs were cultured in serum free E8 media, starvation was carried out using an E8 medium without the E8 supplements (containing growth factors) for 4 hours. iPSCs were stimulated for 10 minutes by changing the media back to the E8 medium containing E8 supplements. Additionally, DLD-1 was treated with pervanadate (PV) for 15 minutes using 0.5 mM and 1mM of PV. A 100mM PV solution was prepared by adding in equal volumes 100 mM sodium orthovanadate and 100 mM 30% H_2_O_2_ in 20 mM HEPES at pH 7.5 ^82^. All cells were washed twice with ice-cold PBS containing phosphatase inhibitors (2mM of sodium orthovanadate, 1mM of beta-glycerophosphate, and 1mM of sodium fluoride). The cells were lysed with the addition of 200 μL of boiling lysis buffer (5% SDS (sodium dodecyl sulfate), 100mM Tris pH 8.5, 5mM TCEP (Tris(2-carboxyethyl)phosphine hydrochloride), and 10mM CAA (Chloroacetamide)). The cells were scraped off the plate, and the resulting lysates were heated for 10 minutes at 95 °C and sonicated using a probe sonicator (Sonic dismembrator, Fisherbrand) with two pulses of 5 seconds at 40% amplitude. Protein concentrations were determined by BCA (bicinchoninic acid) method (Pierce).

### 3. Digestion, desalting

600 μg of protein per sample was digested overnight using protein aggregation capture (PAC)-based digestion ^83^, implemented on a KingFisher Flex platform ^84^. Digestion was carried out on hydroxyl beads (ReSyn Bioscience) at a protein-to-beads ratio of 1:2, with endoproteinase LysC (FUJIFILM Wako Pure Chemical Corporation) and trypsin (Sigma-Aldrich) in an enzyme-to-protein ratio of 1:500 and 1:250, respectively. Samples were then acidified to a final concentration of 1% FA (formic acid). The equivalent of 750ng of peptides was aliquoted for proteome analysis. The resulting peptides were desalted on C_18_ Sep-Pak 96-well plates (40mg sorbent, Waters).

### 4. Phosphopeptide enrichment

Phosphopeptide enrichment was carried out on a KingFisher Flex robot ^84,85^ using Ti-IMAC HP magnetic beads (ReSyn Bioscience) at a peptide-to-beads ratio of 1:2. C18-bound peptides were directly eluted from the Sep-pak to the KingFisher plate using 75 μL of 80% ACN (acetonitrile)^86^ and completed with 200 μL of concentrated loading buffer (80% ACN, 8% TFA, 0.6 M glycolic acid). Briefly, the magnetic beads were added to 500 μL of ACN in plate #2, washed with 500 μL loading buffer (80% ACN, 5% TFA, 0.1M glycolic acid) in plate #3, added to the peptide samples in plate #4, washed in plate #5 (500 μL of loading buffer), plate #6 (500 μL of washing buffer 2: 80% ACN, 5% TFA), plate #7 (500 μL of washing buffer 3: 10% ACN, 0.2% TFA), and eluted in 200 μL 1% ammonia in plate #8. The enriched phosphopeptides were acidified with 40 μL of 10% TFA and loaded on Evotips, according to the manufactureŕs protocol for subsequent LC-MSMS analysis.

### 5. EGF Stimulation and low input analysis

Hela cells for EGF stimulation were cultured until 70% confluence and serum-starved overnight. Cells from two conditions; untreated control and stimulated with 20 nM EGF (Peprotech) for 8 min, were washed twice with ice-cold PBS and lysed and quantified as described above. Proteins from each condition were bulk-digested using PAC and the peptide concentration for each condition was determined using a nanodrop spectrophotometer (NanoDrop 2000C, Thermo FIsher Scientific) at 280 nm. 1, 5, 10 to 20 ug peptides were enriched for phosphopeptides as described above in 5 replicates each condition. The enriched phosphopeptides were acidified with 10% FA and loaded on Evotips, according to the manufactureŕs protocol for subsequent LC-MSMS analysis.

### 6. LC-MS/MS

All samples were analysed on the Evosep One system (Evosep Biosystems, Denmark) coupled to an Orbitrap-Astral mass spectrometer (Thermo Fisher Scientific, Germany). Standard gradients were used with commercial Evosep columns (30 SPD (samples per day) with EV1137: 15 cm x 150 μm, 1.5 μm - 100 SPD with EV1109: 8 cm x 150 μm, 1.5 μm) connected to a stainless steel emitter (30 μm i.d., Evosep, EV1086), with temperature set to 40°C through an externally controlled butterfly heater (MSWil, Cat#PST-ES-BPH-20). The columns were interfaced to the mass spectrometer using an EASY-Spray source. Whisper zoom gradients were used with Aurora Rapid columns (5 cm x 75 μm, 1.7 μm) interfaced to the mass spectrometer using a Nanospray Flex source, with column temperature maintained at 50°C.

The Orbitrap-Astral mass spectrometer was operated in positive mode, with a spray voltage set at 2.0kV (1.8 kV for whisper zoom gradients),funnel RF level at 40, and heated capillary temperature at 275°C. All files were acquired using narrow window data independent acquisition (nDIA) ^21^ with a full scan resolution of 240,000 with a normalized automatic gain control (AGC) target of 500%.

For proteome analysis, the Orbitrap operated across a mass-to-charge ratio range from 380 to 980 m/z with a maximum injection time set at 3 ms, while it operated over a range from 480 to 1080 m/z with a maximum injection time set at 30 ms for phosphoproteomics analysis. Tandem mass spectra were generated across the respective mass range set for MS1 analysis with a scan range from 150 to 2,000 m/z. Peptide fragmentation was carried out using higher-energy collisional dissociation with a normalised collision energy set at 25%. The normalised AGC target was set at 500 %. Different methods with specific isolation window sizes and maximum injection times were used based on the chromatographic gradient employed. 30 SPD was used with 2Th isolation windows and 2.5 ms maximum injection time, 100 SPD was used with 4Th isolation windows and 2.5 ms maximum injection time, 80 SPD was used with 4Th isolation windows and 6 ms maximum injection time, and 120 SPD was used with 4Th isolation windows and either 3ms or 6 ms maximum injection time.

### 7. Data searching

#### 7.1. Spectronaut

All raw files were searched in Spectronaut version 19 (Biognosys) against the human protein reference database from Uniprot (2022, 20,598 entries) supplemented with a contaminant database from Spectronaut (246 entries). Each cell line was searched separately, and the resulting SNEs were combined using the “combine SNE” function. Libraries were generated from the search archives, therefore using the parameters set in the Pulsar search.

Cysteine carbamidomethylation was set as a fixed modification, and N-terminal acetylation and methionine oxidation were set as variable modifications, with a maximum number of variable modifications kept at 5, and a maximum number of missed cleavages kept at 2. For phosphoproteomics, serine, threonine, and tyrosine phosphorylation were added as variable modifications. Cross-run normalization was allowed, with a modification filter on the phosphoproteomics files, where only phosphorylated peptides were used for normalization. The maximum and minimum number of fragments per peptide were set to 25 and 6, respectively. Fragment ion selection was set to evidence-based.

For phosphoproteomics analysis, the PTM localisation filter was checked, with a threshold of 0.75. Library-based searches were performed using the same parameters as directDIA.

#### 7.2. DIA NN

All raw files were searched on DIA-NN 2-1. In silico phosphopeptide spectral library was generated using DIANN with the human reference database from Uniprot (2022, 20,598 entries) supplemented with a contaminant database from Spectronaut (246 entries). 2 missed cleavages were allowed for a maximum of 3 variable modifications per precursor, including N-terminal methionine excision, cysteine carbamidomethylation, and serine, threonine, tyrosine phosphorylation. The peptide length range and precursor charge range were kept to default, while the precursor and fragment mass ranges were adapted to the MS method parameters: 480 to 1080 m/z for the precursors and 150 to 2000 m/z for the fragments.

Empirical spectral library-based and in silico spectral library-based searches were carried out in DIANN using the above-mentioned parameters. Proteoform scoring was used for phosphoproteomics data.

Fragmentation spectra from the experimental library generated with Spectronaut were compared to the fragmentation profiles obtained from the in silico predicted spectral library using DIA-NN. The cosine correlation spectral angle was measured per phosphoprecursor between the intensities of equivalent fragments. Cosine correlation was calculated using ‘matchms’^87,88^ library in python

### 8. Data analysis

#### 8.1. Data processing

The phosphosite table was exported from Spectronaut, using Spectronaut’s default filtering parameters, filtered for localisation probability > 0.75, and entries without gene information were filtered out. Cell lines were split into different tables and transformed into PhosphoExperiment objects (ppe) using the PhosR package ^89^. Each cell line was filtered for valid value, allowing to conserve phosphosites quantified in 3 out of 4 replicates in at least one condition (Ctrl and/or Stim) before proceeding with two steps imputation: 1) site- and condition-specific imputation, imputing sites with ≥ 50% valid value in one condition, sampled from the empirical normal distribution from the same condition, followed by 2) tail-based imputation. All cell lines were joined into a unique table and normalized through mean centering. Fold changes were calculated per site and per cell line against a background composed of the mean intensity across all cell lines. If a site was unique to a cell line, a positive fold change of 1.5 was applied.

Proteome data was analyzed on the protein group level, each cell line was splited into different tables and filtered for valid values, allowing protein groups with at least 3 valid values out of 4 in at least one condition. No imputation step was carried out. The tables were joined into a unique table and all intensities were normalized through mean centering. Fold changes were calculated per protein group per cell line against a background composed of the mean intensity across all cell lines.

#### 8.2. DIANN counts and filtering

For benchmarking DIANN and Spectronaut on differentiated stem cells, DIANŃs outputs were processed to match Spectronaut’s. Phosphosites were defined with a localisation probability ≥ 0.75 and included multiplicity.

The Parquet report was opened, and entries without gene information were filtered out. The following filters were applied: PG.Q.Value ≤ 0.05, Global.PG.Q.Value ≤ 0.01, Lib.PG.Q.Value ≤ 0.01 (when match between runs (MBR) was on), Quantity.Quality ≥ 0.5 and PG.MaxLFQ.Quality ≥ 0.7 according to the software’s guidelines. The table was further filtered to only keep phosphopeptides. Phosphopeptides were annotated and collapsed per site and per multiplicity before being filtered based on localisation probability with a cutoff at 0.75.

#### 8.3. Stoichiometry calculation

Site stoichiometry was calculated from the raw phosphoproteomics and raw corresponding proteomics datasets (i.e., non-filtered and non-normalised), following the approach of Olsen et al ^19^. Precursor intensities were first collapsed to obtain peptide-sequence–level intensities. Phosphopeptide precursor intensities were then grouped by peptide sequence and by the position of the phosphorylated residue within the protein sequence. Phosphoproteomics and proteomics tables were merged based on protein groups and peptide sequence, resulting in protein-, peptide-, and site-sequence–level intensities required for stoichiometry estimation.

Data from the serum-stimulated and control conditions were pre-processed separately. Phosphorylation site-sequences with fewer than two valid values per cell line were removed in each dataset independently. For each phosphorylation site-sequence, the median intensity and standard deviation were computed in both conditions, and the two tables were subsequently merged. Entries lacking median intensities in either the phosphoproteome or proteome were discarded. To retain only sites exhibiting measurable changes, we kept rows where the median intensity in the stimulated condition fell outside the interval defined by the control median ± 3 × control standard deviation.

Three ratios (x, y, and z) were calculated as the non-log-transformed stimulated/control intensity ratios at the site/peptide-sequence, peptide-sequence, and protein levels, respectively. As no global protein abundance changes were expected, entries with protein-level ratios outside the range 0.8 < z < 1.2 were removed.The following proportions of phosphopeptide to non-phosphopeptide were calculated:

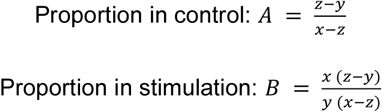

Stoichiometries were then calculated as A/(A+1) or B/(B+1), depending on the condition.Further filtering of the data was required. When no regulation occurred, x and y ratios were nearly identical (ie. abs(x-y) < 0.01), which produced extreme stoichiometry values. These cases were labelled as “not changing” and removed. Additionally, A and B must be positive to yield valid stoichiometries, as this assumes that the total amount of a given peptide (phosphorylated plus non-modified) remains constant between conditions. Entries where this assumption was not respected were labelled as “breaking the assumption” and excluded.

#### 8.4. Kinase score calculation

##### 8.4.1. Kinase activity scores

Unfiltered in-vitro kinase reactions relationships were downloaded ^32,64^ and reprocessed to avoid the first filtering applied in the iKiP-DB. Kinase-substrate relationships were combined with PhosphoSite Plus annotation and filtered to exclude phosphosites that were substrates for over 20 kinases. Kinases with fewer than 5 substrates were filtered out. The modified iKiP-DB contains kinase substrate relationships for 369 kinases.

Kinase activity inference was carried out using the modified iKiP database with the PTM-SEA algorithm for ssGSEA ^31^. The default parameters were used (ie. sample normalisation type as rank, weight at 0.75, statistics using “area under RES”, the output score coming from the normalised enrichment score, number of permutation set at 1000, correlation type being Z-score) except for the minimum overlap, which was decreased to 5.

Motif enrichment analysis was carried out using the Kinase Library ^62^. Serine/Threonine and Tyrosine were processed separately for each cell line. For carrying out multiple comparisons, the publicly available code of the Kinase Library was used in Python. Default parameters were used (ie. percentile rank as enrichment metric, an enrichment threshold of 15 for S/T and 8 for Y).

Activation loop sites were identified based on an activation loop database from the Phomics algorithm. Briefly, activation loops were identified in kinases present in the human proteome database based on their amino acid sequence. The activation loop domain is defined between “DFG” and “APE” amino acid sequences, or homologues. All serine, threonine, and tyrosine amino acids in those loops were compiled and complemented with known activation sites from PhosphoSite Plus to generate an “activation” database containing activation sites over 327 kinases.

Kinase activity were inferred using Rokai using the combined PSP and Signor kinase substrate dataset, with Zscores used as a metric.

##### 8.4.2. Cscore calculation

A kinase activity score was compiled using the proteome and phosphoproteome datasets (Cscore). All calculations were performed on log2 fold changes of one cell line against the mean of all cell lines.

The Cscore is composed of two entities combined as a geometric mean. The first term represents the evidence of the presence of the kinase at the proteome and phosphoproteome levels, while the second term represents the evidence of kinase activity on the phosphoproteome level.

The first score describes the kinase intensity in the proteome data. The score originates from the log2 fold change of the intensity of the kinase in a given cell line and the average intensity of that given kinase across all cell lines. The second score represents the intensity of the kinase on the phosphoproteome. The score was computed as an enrichment type score, where the mean log2 fold changes of all sites on a given kinase were subtracted to the mean log2 fold changes of all sites in the cell line. The difference was then multiplied by the square root of the number of sites quantified in the kinase.

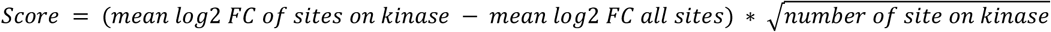

Two substrate activity scores were computed. The first one takes into account kinase substrate relationships based on in vitro kinase reactions established through the use of a modified iKiP database and the PTM SEA algorithm. The resulting score originated from the product of the resulting normalised enrichment score (NES) with the -log10 FDR. Likewise, the second substrate activity score is based on motif enrichment analysis using the Kinase Library. The resulting score originates from the product of the NES with the -log10 FDR. Finally, the activation sites score was computed as an enrichment type score, where the log2 fold change of all activated sites was subtracted with the log2 fold change of all sites, and subsequently multiplied by the square root of the number of activation sites quantified.

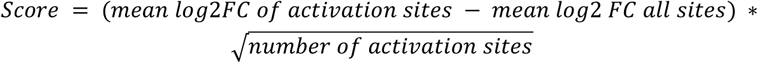

All five scores underwent sigmoid transformation to scale them from 0 to 1 without compressing the scores.

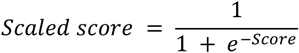

Finally, all scores were combined as follows:

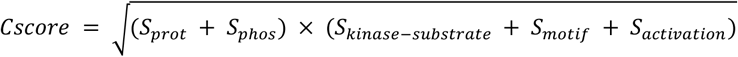

Cell line–specific vulnerabilities were identified with the calculation of a Z-score for each kinase across all cell lines. Kinase–cell line pairs with a Z-score >2 (i.e., activity > median + 2 standard deviations) were considered outliers, while kinases with a kinase activity score >0.5 detected in four or fewer cell lines were classified as unique to those lines.

## Data availability

The mass spectrometry proteomics data have been deposited to the ProteomeXchange Consortium (http://proteomecentral.proteomexchange.org) via the PRIDE ^90^ partner repository with the data set identifier PXD068583.

## Author contribution

C.K. and J.V.O. designed the proteomics experiments, C.K., H.C., K.B.E., I.P., P.S. and S.L.J. prepared and performed proteomics experiments and analyzed the resulting data. A.M.V and J.V.O. critically evaluated the results. C.K. wrote the first draft of the paper. All authors read, edited and approved the final version of the paper.

## Acknowledgements

We would like to acknowledge the following people for providing cell stocks, dishes, or facilitating our access to cell lines for the project: Maico Lechner, Katrin Stuber, Leander Van der Hoven, Holda Anagho-Mattanovich, Anders Handrup Kverneland, Florian Mau, and Juliette Ferrand.

## SUPPLEMENTARY FIGURES

**Supplementary figure 1:**
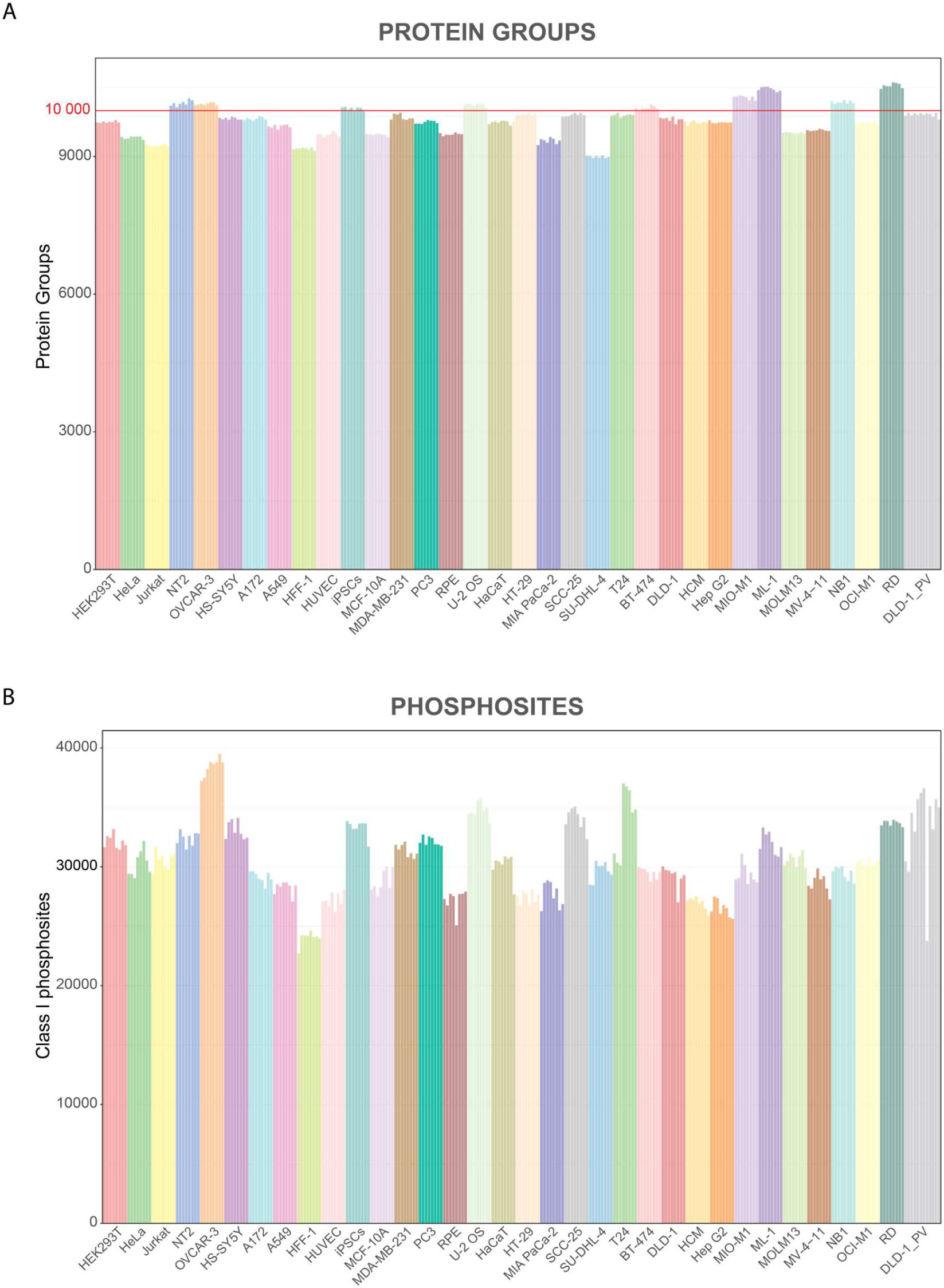
Protein group and phosphosite counts. A- Protein group count per raw file after filtering for valid values within a cell line (¾ valid values in at least one condition (ie. control or serum-stimulation)). B- Class 1 phosphosite count per raw file after filtering for valid values within a cell line (¾ valid values in at least one condition (ie. control or serum-stimulation)).

**Supplementary figure 2:**
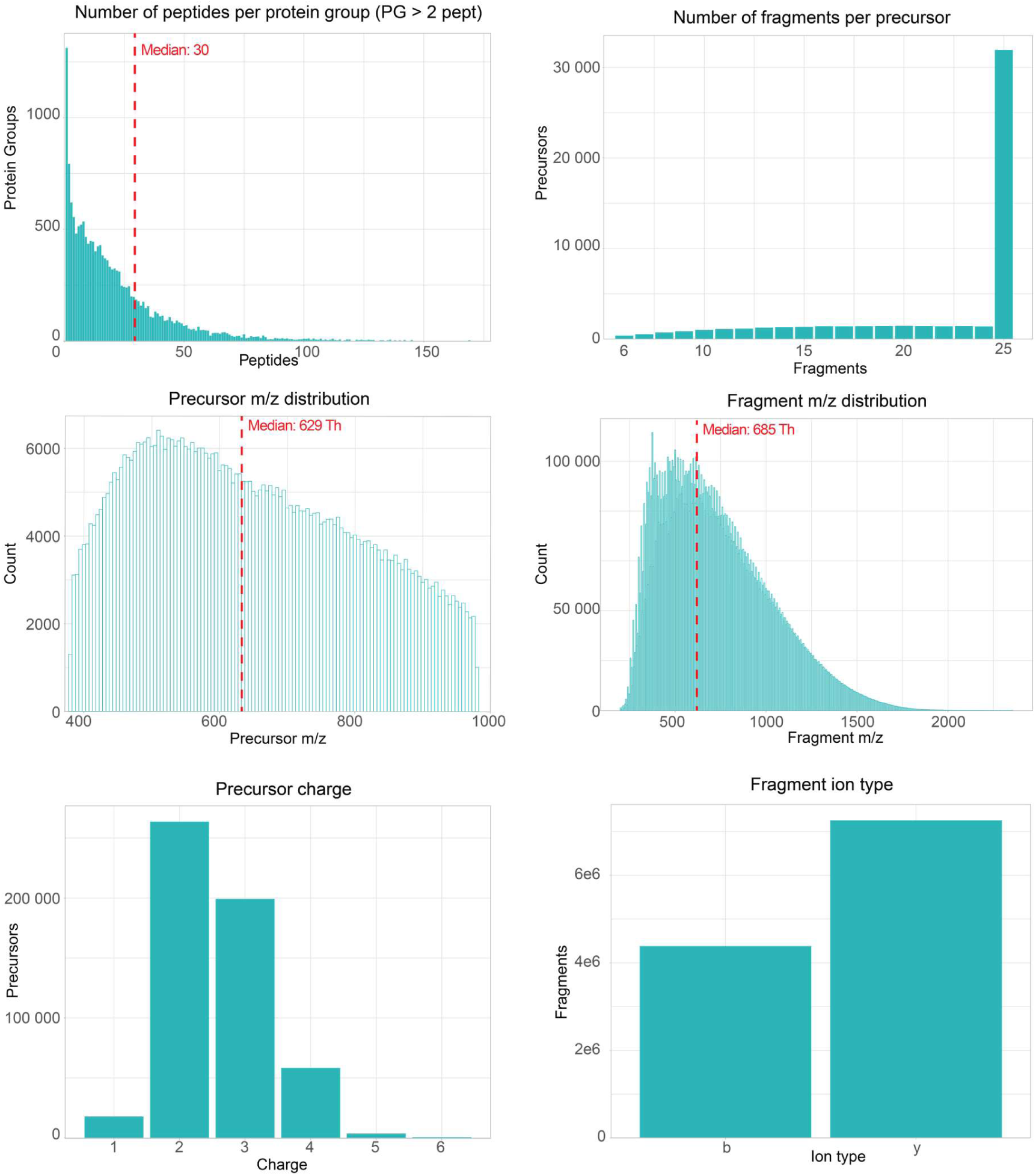
Characteristics of the peptide spectral library. Distributions of the number of peptides per protein groups, number of fragments per precursors, precursor and fragments mass-to-charge ratios, charge states and fragment ion type.

**Supplementary figure 3:**
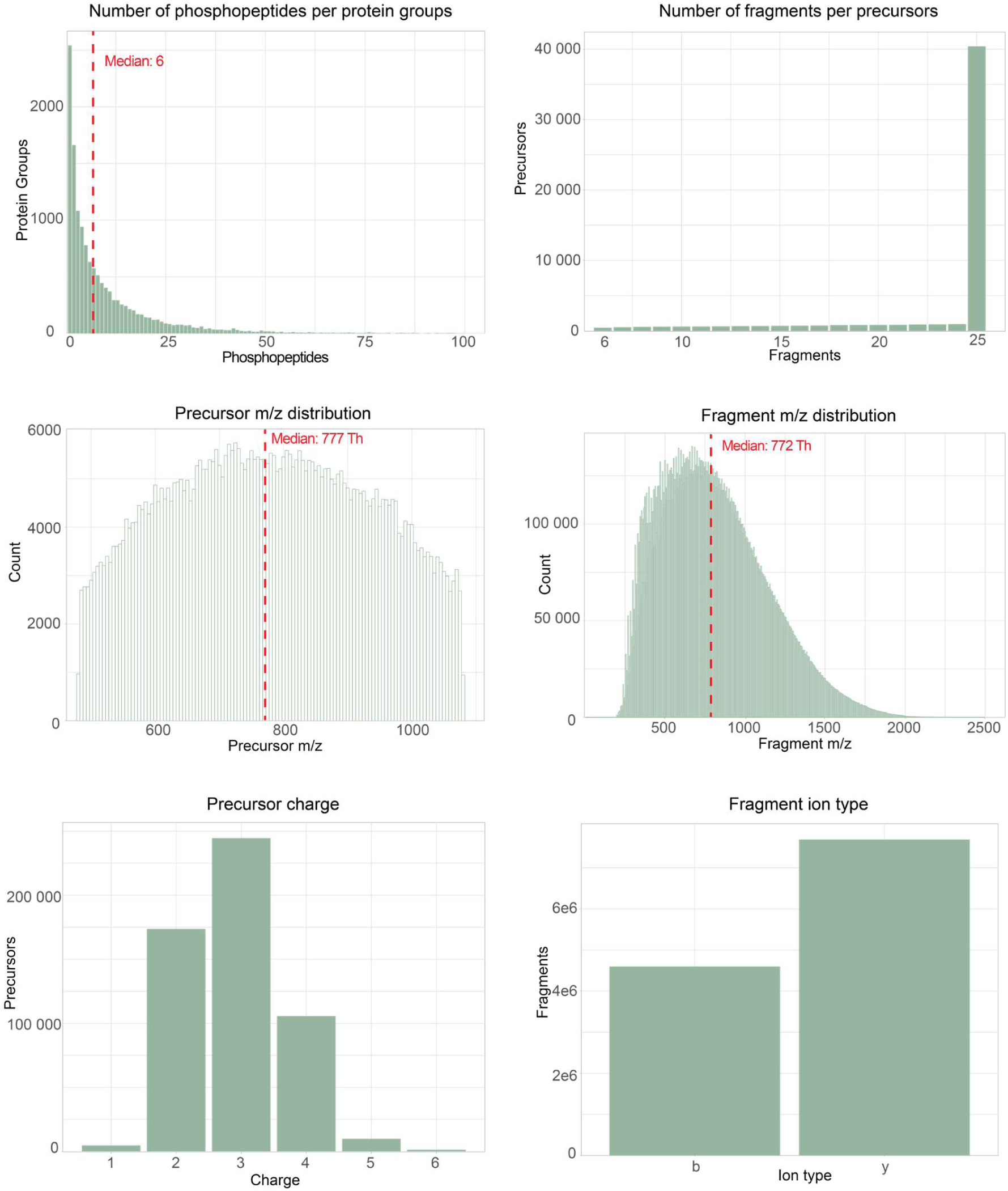
Characteristics of the phosphopeptide spectral library. Distributions of the number of phosphopeptides per protein groups, number of fragments per precursors, precursor and fragments mass-to-charge ratios, charge states and fragment ion type.

**Supplementary figure 4:**
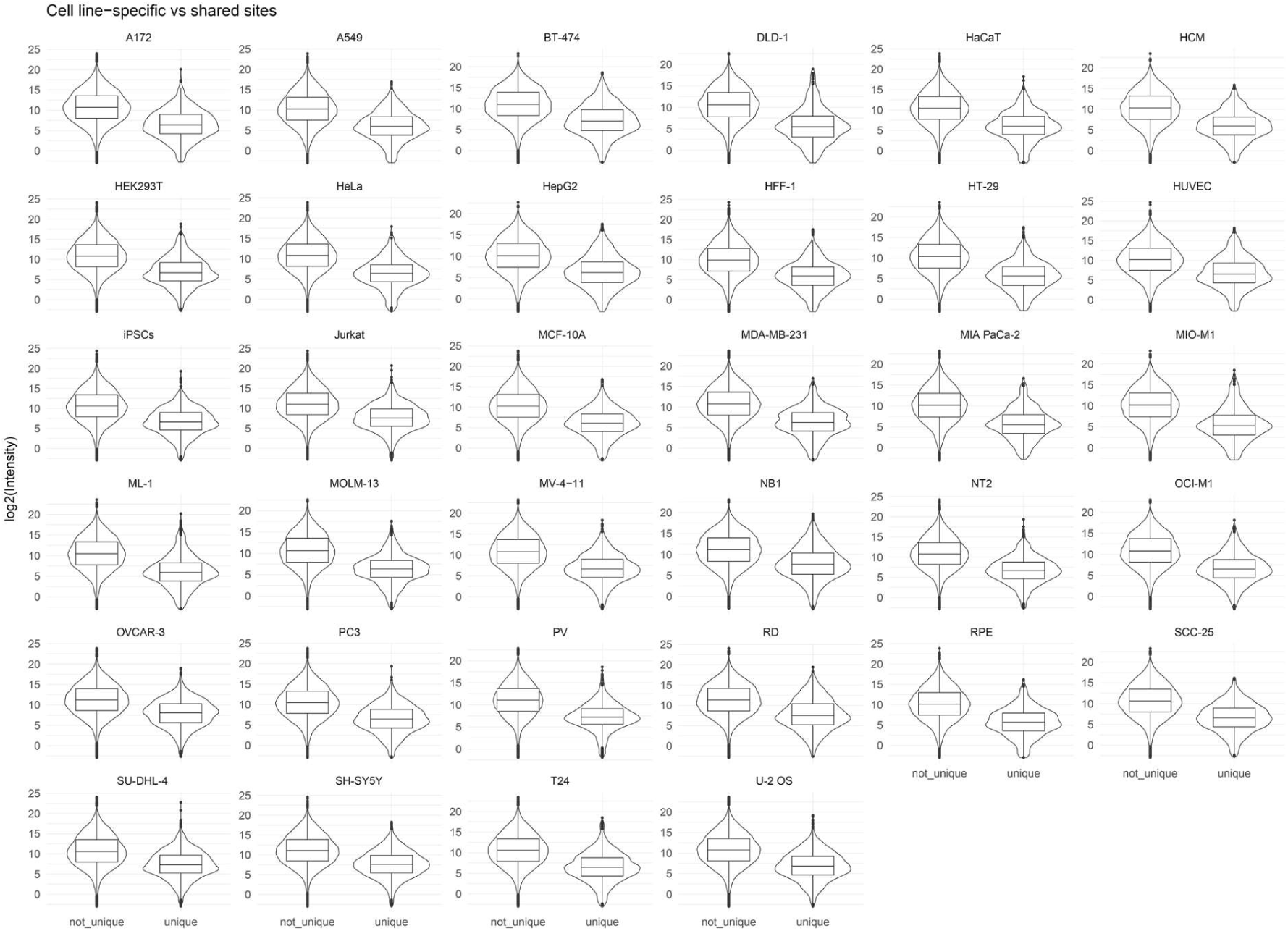
**Intensity distribution between cell-line specific phosphosites and non-cell line specific ones.**

**Supplementary figure 5:**
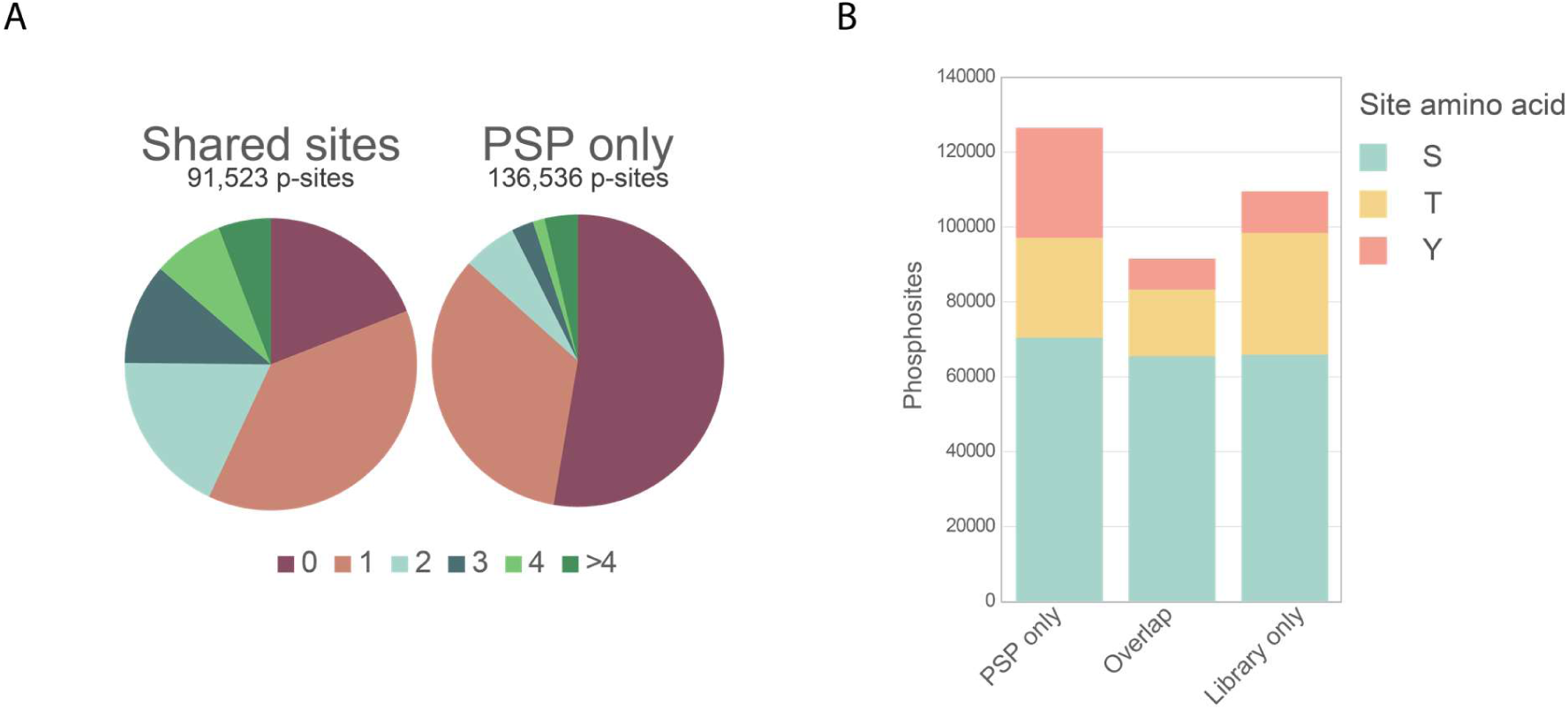
Comparison of phosphosites from PSP and the generated library. A-Venn diagrams showing the proportion of sites with 0, 1, 2, 3, 4 or above 4 references in the literature using MS-based techniques in the overlap between the library and PSP, and in the sites unique to PSP. B- Phosphosite count overlapping between the library and PSP, unique to PSP and unique to the library, with their respective proportion of pS, pT and pY.

**Supplementary figure 6:**
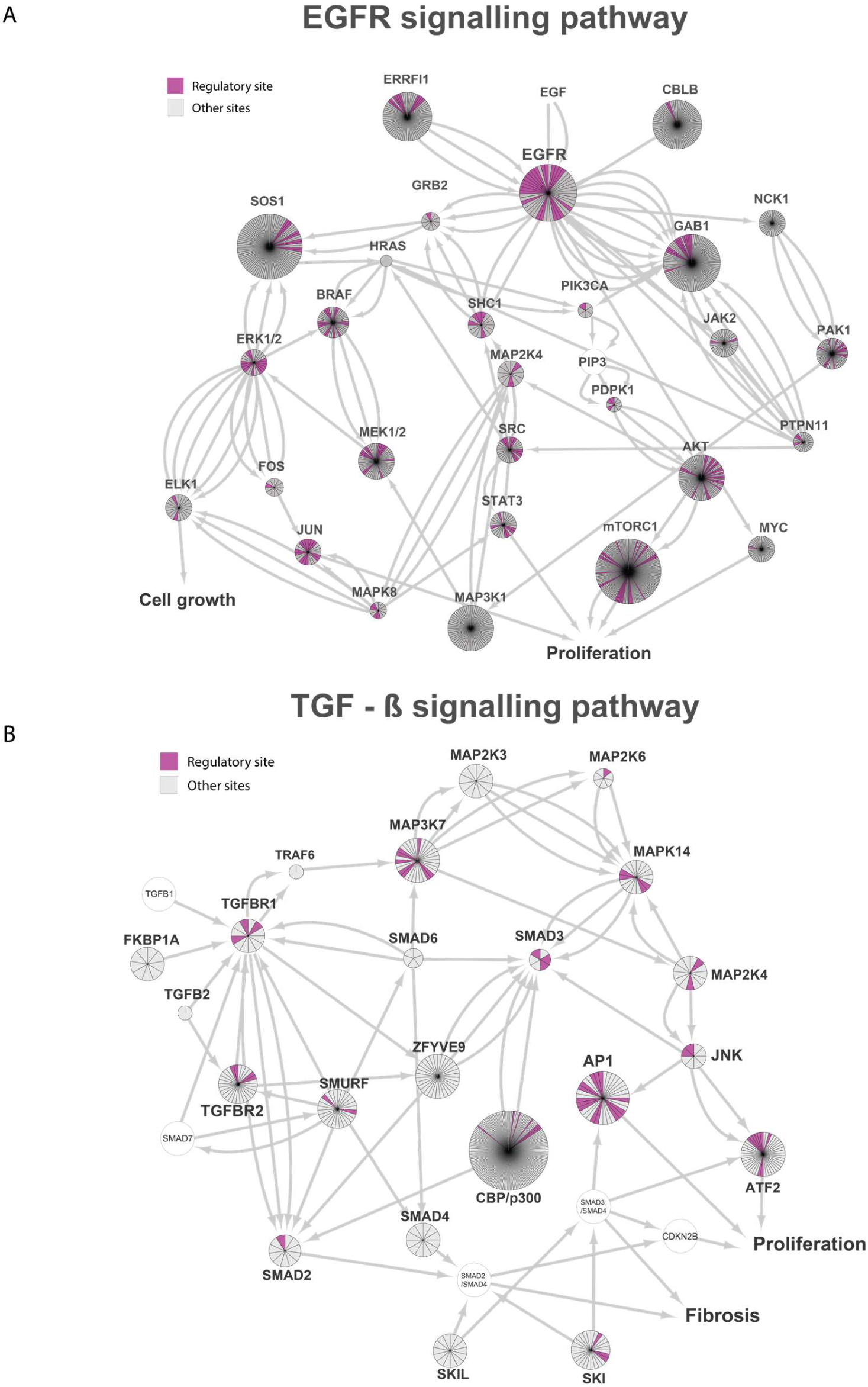
Phosphosite coverage on signaling pathways. A- Phosphosite coverage of the EGFR signaling pathway represented as a network. Each slice on the genes represents a phosphosite covered in the library on the given gene. Pink sites are known regulatory sites from PSP. B- Phosphosite coverage of the TGF-beta signaling pathway represented as a network. Each slice on the genes represents a phosphosite covered in the library on the given gene. Pink sites are known regulatory sites from PSP.

**Supplementary figure 7:**
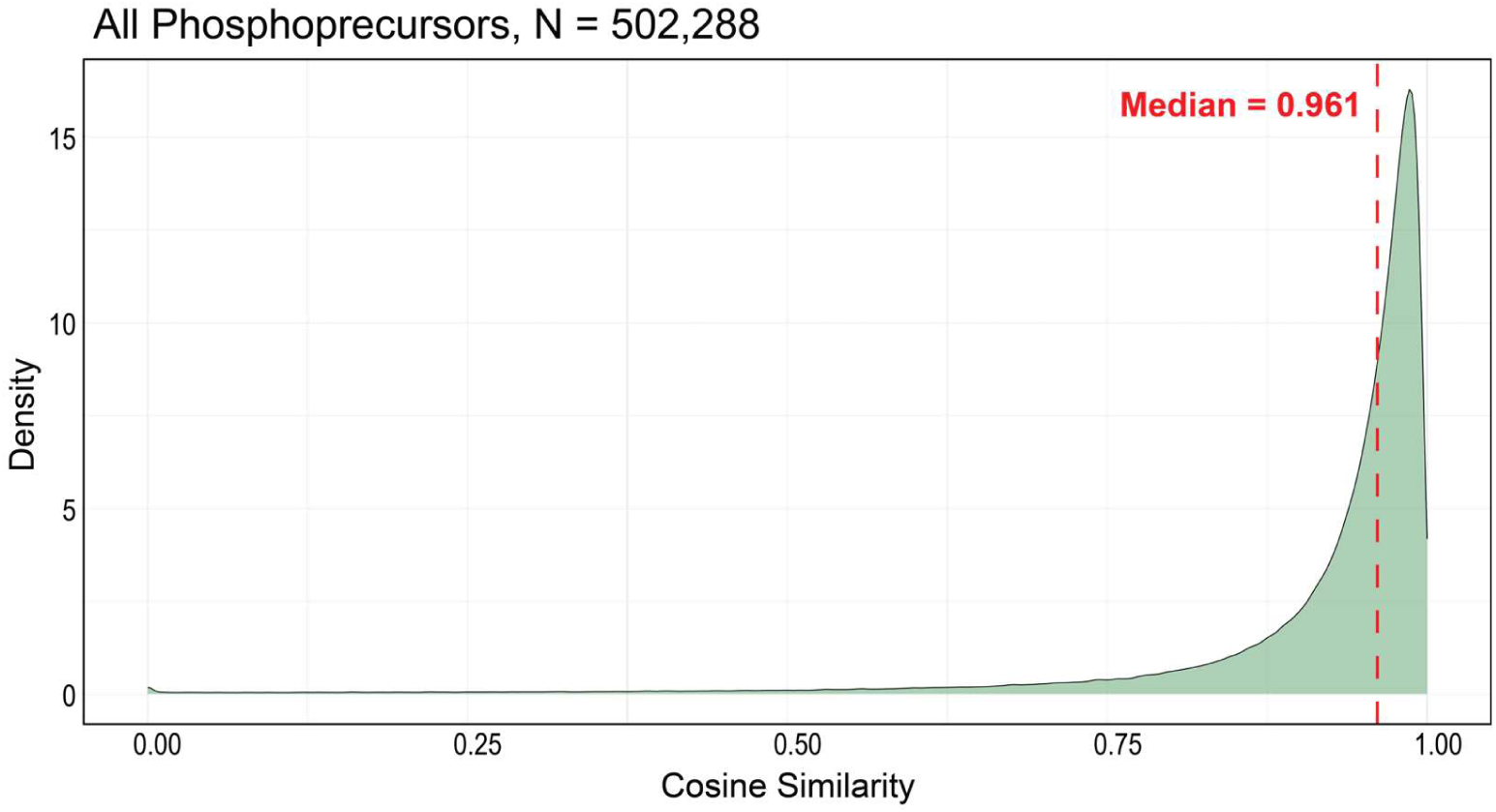
Empirical and in silico generated spectral phosphopeptide libraries correlation. Density plot displaying the cosine correlation spectral angle between the intensities of fragment pairs per precursor between both spectral libraries. The median spectral angle across all 502,288 phosphoprecursors is displayed in red.

**Supplementary figure 8:**
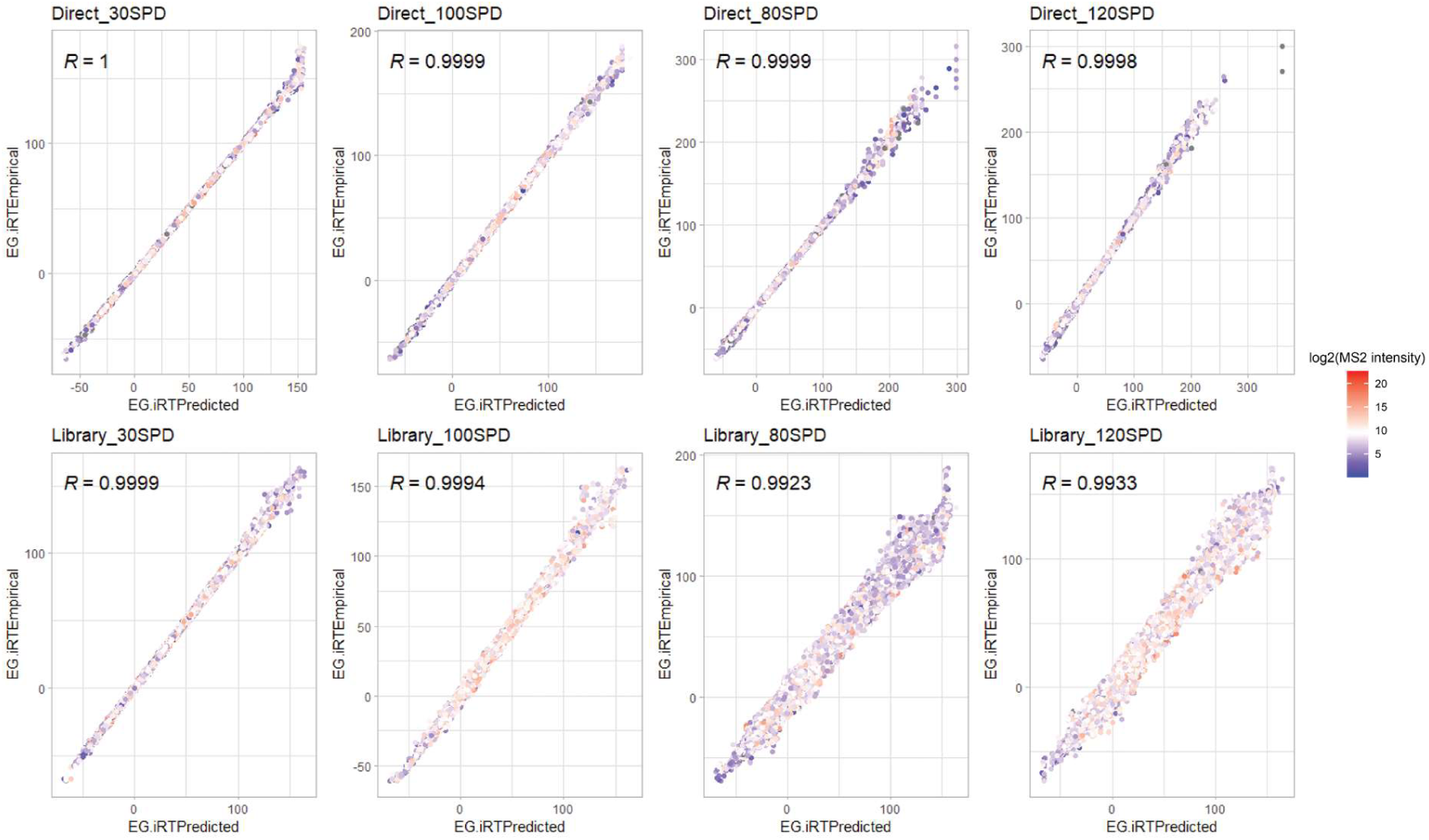
Retention time correlation between measured retention times and predicted retention times. In the case of directDIA, the predicted retention times are generated during the pulsar search on the data itself. In the library-based approach, the predicted retention times are indexed retention times from the library, which was generated using 30 SPD. Spearman correlation coefficients were calculated and are displayed on the plot. The color represents the log2 MS2 intensity.

**Supplementary figure 9:**
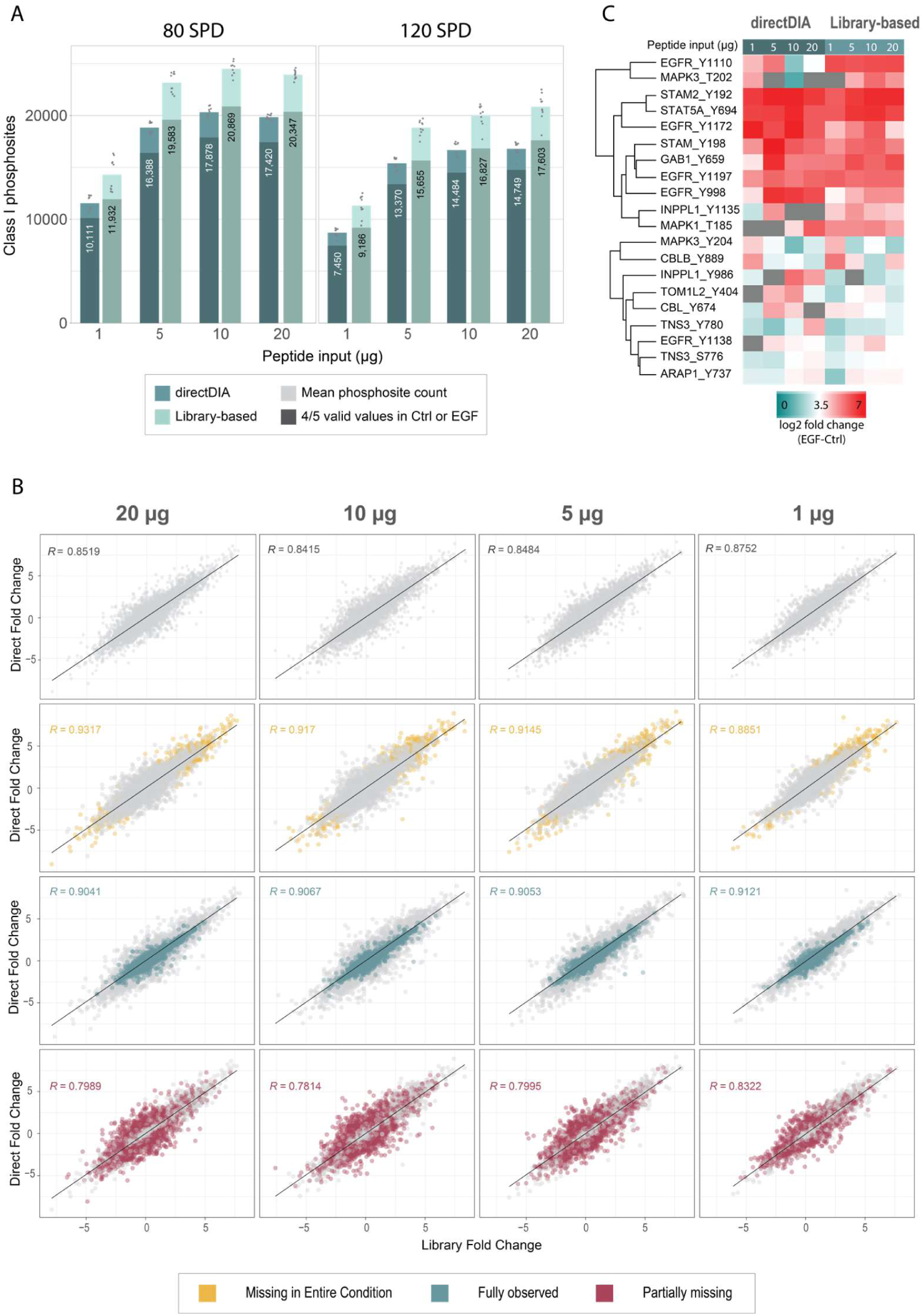
Performance of the spectral library approach compared to directDIA in a scale down. A-. Class 1 phosphosite coverage on peptide input amount ranging from 1 to 20 μg, using the 80 SPD and the 120 SPD gradients. The dots represent the site count per replicate. The lightest bar displays the mean site count across the 5 replicates in the control and EGF stimulated conditions (N=10), and the darkest bar shows the site count after valid value filtering for ⅘ valid values in the control or stimulated experiments. The number displayed corresponds to the site count after filtering. B- Heatmap of the main upregulated phosphosites in the EGFR canonical pathway. C-Correlation between the calculated fold changes between the EGF stimulated condition and the control, in directDIA and using the empirical spectral library. Spearman correlation coefficient is calculated per input amount and per phosphosite population: Missing in entire condition, fully observed or partially missing.

**Supplementary figure 10:**
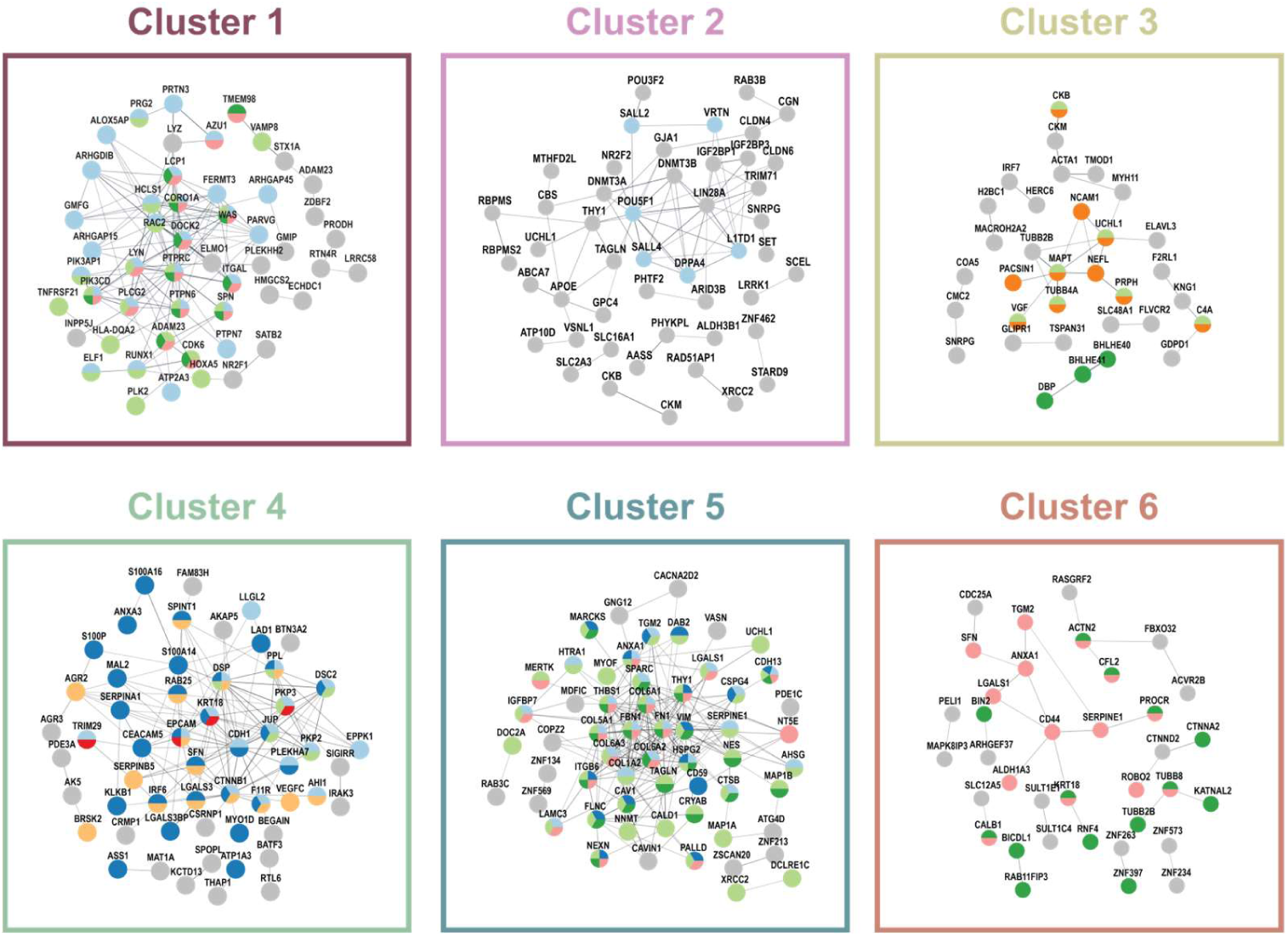
Functional protein association network of the top 100 most differentially expressed proteins per cluster. The plot was generated using Cytoscape and the StringApp ^46^. Singletons are not displayed. The color of each protein displays which protein was used per enriched gene set in Figure 5C.

**Supplementary figure 11:**
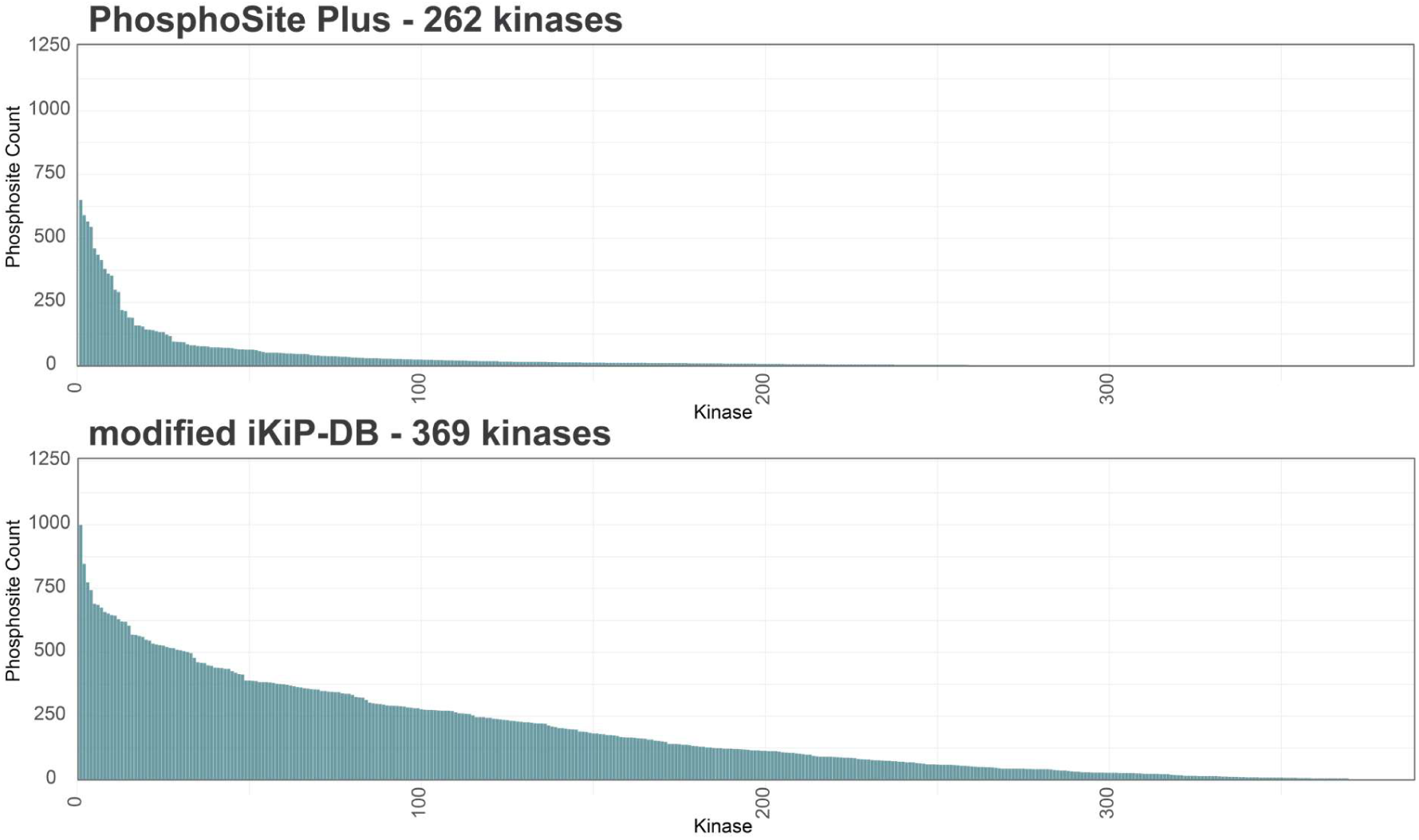
Overview of the number of kinase-substate relationships in the PSP and modified iKiP-DB databases. Kinases are ranked based on their number of substates in the respective database.

**Supplementary figure 12:**
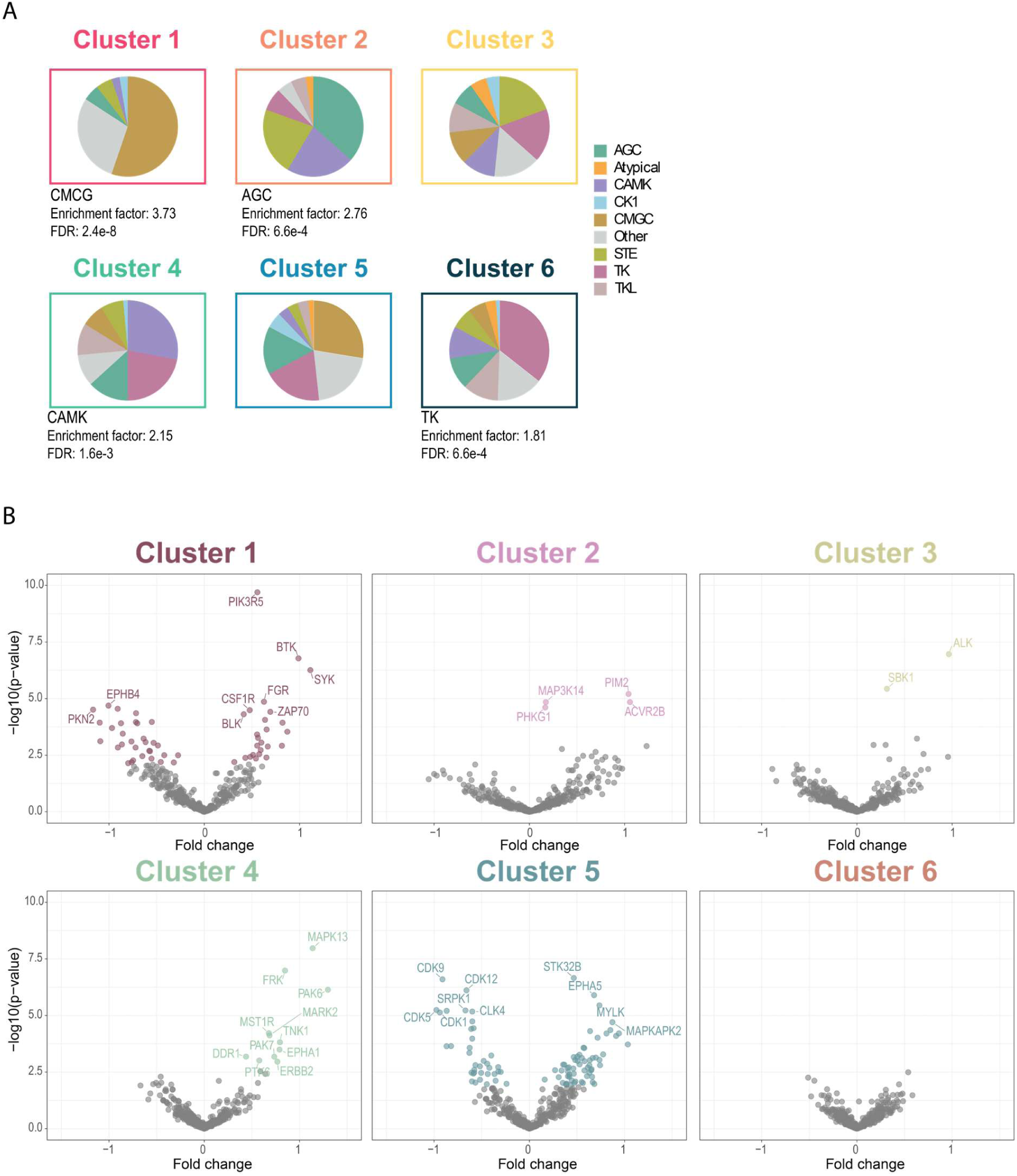
Combined kinase score allows to study kinase family enrichments and specific activity changes across functional cell line clusters. A- Distribution of the kinase families per cluster, which were defined Figure 6C. Fisher enrichment analysis was carried out for each cluster. Benjamini-Hochberg correction was applied for FDR calculation. B- Volcano plot for each functional cell line cluster, which were defined Figure 5B. Fold changes were calculated per cluster against the background of all the other cell lines. Significance was assessed using Student’s t-test and colored dots represent kinases with an activity statistically significantly changed at a FDR of 5%.

